# Cysteine depletion triggers adipose tissue thermogenesis and weight-loss

**DOI:** 10.1101/2024.08.06.606880

**Authors:** Aileen H. Lee, Lucie Orliaguet, Yun-Hee Youm, Rae Maeda, Tamara Dlugos, Yuanjiu Lei, Daniel Coman, Irina Shchukina, Sairam Andhey, Steven R. Smith, Eric Ravussin, Krisztian Stadler, Fahmeed Hyder, Maxim N. Artyomov, Yuki Sugiura, Vishwa Deep Dixit

**Affiliations:** Department of Pathology, Yale School of Medicine, New Haven, CT 06520, USA; Department of Comparative Medicine, Yale School of Medicine, New Haven, CT 06520, USA; Department of Immunobiology, Yale School of Medicine, New Haven, CT 06520, USA; University of Kyoto, Japan; Department of Radiology and Biomedical Imaging, Yale University; Department of Biomedical Engineering, School of Engineering and Applied Science, Yale University; Department of Pathology and Immunology Washington University School of Medicine, St. Louis, MO 63110, USA; Translational Research Institute for Metabolism and Diabetes, AdventHealth, Orlando, FL, USA; Pennington Biomedical Research Center, Baton Rouge, LA, USA; Yale Center for Research on Aging, Yale School of Medicine, New Haven, CT 06520, USA

## Abstract

Dietary interventions such as caloric restriction (CR)^1^ and methionine restriction^2^ that prolong lifespan induce the ‘browning’ of white adipose tissue (WAT), an adaptive metabolic response that increases heat production to maintain health^3,4^. However, how diet influences adipose browning and metabolic health is unclear. Here, we identified that weight-loss induced by CR in humans^5^ reduces cysteine concentration in WAT suggesting depletion of this amino-acid may be involved in metabolic benefits of CR. To investigate the role of cysteine on organismal metabolism, we created a cysteine-deficiency mouse model in which dietary cysteine was eliminated and cystathionine γ-lyase (CTH)^6^, the enzyme that synthesizes cysteine was conditionally deleted. Using this animal model, we found that systemic cysteine-depletion causes drastic weight-loss with increased fat utilization and browning of adipose tissue. The restoration of dietary cysteine in cysteine-deficient mice rescued weight loss together with reversal of adipose browning and increased food-intake in an on-demand fashion. Mechanistically, cysteine deficiency induced browning and weight loss is dependent on sympathetic nervous system derived noradrenaline signaling via β3-adrenergic-receptors and does not require UCP1. Therapeutically, in high-fat diet fed obese mice, one week of cysteine-deficiency caused 30% weight-loss and reversed inflammation. These findings thus establish that cysteine is essential for organismal metabolism as removal of cysteine in the host triggers adipose browning and rapid weight loss.

## Main

The <u>C</u>omprehensive <u>A</u>ssessment of <u>L</u>ong-term <u>E</u>ffects of <u>R</u>educing <u>I</u>ntake of <u>E</u>nergy (CALERIE-II)clinical trial in healthy adults demonstrated that a simple 14% reduction of calories for two years without any specific dietary prescription to alter macronutrient intake or meal timings can reprogram the immunometabolic axis to promote healthspan^5,7,8^. Harnessing the pathways engaged by CR in humans may expand the current armament of therapeutics against metabolic and immune dysfunction. Induction of negative energy balance and resultant activation of mitochondrial fatty acid oxidation by CR is thought to underlie some of its beneficial effects on healthspan^5^. However, it has also been suggested that CR-induced metabolic effects may be due to decreased protein intake in food-restricted animal models^9,10^. Adding back individual amino acids to calorie-restricted *Drosophila* abolished the longevity effects, and traced to the limitation of methionine, an important node for lifespan extension^10^. Indeed, methionine restriction (MR) in rodents increases lifespan^11^ with enhanced insulin sensitivity, adipose tissue thermogenesis, and mitochondrial fatty acid oxidation^2^. Surprisingly, in long-lived *Drosophila* fed an MR diet, adding back methionine did not rescue the pro-longevity effect of diet, and it was hypothesized that activation of the methionine cycle may impact longevity^10^. Commercial MR diets contain 0.17% methionine compared to normal levels of 0.86%, but notably, the MR diets also lack cystine^12, 13^, another sulfur-containing amino acid (SAA), which is a key substrate for protein synthesis, including synthesis of glutathione, taurine and iron-sulfur clusters^6,14^. Interestingly, in rats, MR-induced anti-adiposity and pro-metabolic effects, including reduction of leptin, insulin, IGF1, and elevation of adiponectin, were reversed when animals were supplemented with cysteine in the diet^15^. Furthermore, cysteine supplementation in MR rats did not restore low methionine, suggesting no increase in the methionine cycle^15^, where homocysteine is converted into methionine via the enzyme betaine-homocysteine S-methyltransferase (BHMT)^6^. The existence of transsulfuration (TSP) in mammals indicates that in case of dietary cysteine scarcity, the host shuttles homocysteine from the methionine cycle via the production of cystathionine, which is then hydrolyzed into cysteine by the enzyme cystathionine γ-lyase (CTH)^6,16^. Cysteine is an ancient molecule that evolved to allow early life to transition from anoxic hydrothermal vents into oxidizing cooler environment^17,18^. Thus, cysteine, the only thiol-containing proteinogenic amino acid, is essential for disulfide bond formation, and redox signaling, including nucleophilic catalysis^6,16^. It remains unclear if cysteine specifically controls organismal metabolism and whether sustained CR in healthy humans can help understand the fundamental relationship between energy balance and sulfur-containing amino acid homeostasis pathways that converge to improve healthspan and lifespan.

### CR in humans reduces adipose tissue cysteine

Adipose tissue regulates organismal metabolism by orchestrating inter-organ communication required for healthy longevity. To study the mechanisms that drive CR’s beneficial effects on human metabolism, we conducted an unbiased metabolomics analysis of the subcutaneous adipose tissue (SFAT) of participants in the CALERIE-II trial at baseline and one year after 15% achieved CR and weight loss^5,7, 8^. The PLSDA analyses of abdominal SFAT biopsies revealed that one year of mild sustained CR significantly altered the adipose tissue metabolome (Fig. 1a). The unbiased metabolite sets enrichment analyses demonstrated significant increases in cysteine, methionine, and taurine metabolism, which indicates rewiring of cysteine metabolism that involves transsulfuration pathway (TSP) (Fig. 1b, c). To investigate the role of TSP in human CR, we re-analyzed our previously reported RNA sequencing data of humans that underwent CR^5,7^. These analyses revealed that compared to baseline, one and two years of CR in humans increased the adipose expression of *CTH* (Fig. 1d) with a concomitant reduction in the expression of *BHMT* (Fig. 1e) suggesting reduction in methionine cycle and shift towards TSP (Fig. 1c). Interestingly, prior studies have found that long-lived rodents upregulate metabolites in TSP that generates cysteine from methionine^19,20^. Consistent with our findings in human CR, data from multiple lifespan-extending interventions in rodents identified upregulation of CTH as a common signature or potential biomarker of longevity^21^.

**Figure 1:**
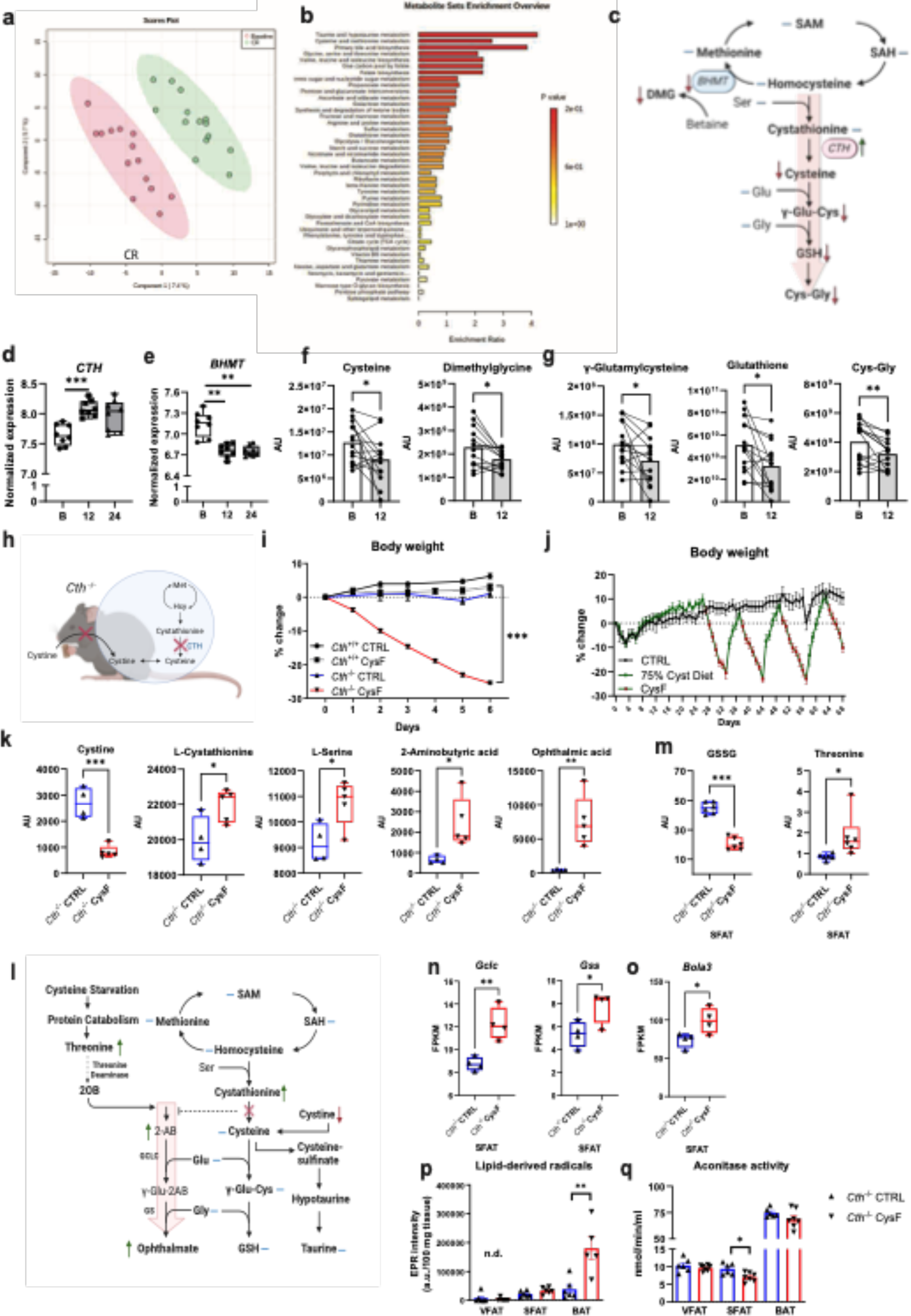
Cysteine deficiency induces weight-loss. a) Principal component analysis of the metabolome of subcutaneous adipose depots (SFAT) of healthy individuals at baseline and after 12 months of caloric restriction (CR) (n=14). b) Metabolite set enrichment analysis shows that compared to baseline, one year of CR in humans activates TSP, with increased cysteine and taurine metabolism. c) Schematic summary of TSP and metabolites from baseline to one year CR, measured in human SFAT. Blue lines indicate unchanged metabolites, green and red arrows indicate significantly increased or decreased metabolites or genes respectively, via paired t-test (p<0.05). d-e) Normalized expression of changes in *CTH*, and *BHMT* in human SFAT at baseline, after 12 months, and 24 months of CR. Adjusted p-values were calculated in the differential gene expression analysis in a separate cohort from metabolome analyses in the CALERIE-II trial (n=8). f-g) Change in metabolites in human SFAT at baseline (B) and 12 months of CR. Significance was calculated using paired t-tests (n=14). AU: arbitrary unit. h) Mouse model used to achieve cysteine deficiency utilizing *Cth*^−/−^ mice fed a Cystine free (CysF) diet. i) Male *Cth*^+/+^ and *Cth*^−/−^ mice were fed control (CTRL) or CysF diets for 6 days (n=5 *Cth*^+/+^ CTRL, n=12 *Cth*^+/+^ CysF, n=8 *Cth*^−/−^ CTRL, n=17 *Cth*^−/−^ CysF, 3 experiments pooled). Percent body weight represented over 6 days of diet. j) *Cth*^−/−^ mice were fed purified control diet (black line) or a diet containing 75% cysteine (green line) alternately switched to CysF diet (green line with red dots n = 6/group). k) Box plots of metabolites involved in TSP in the serum of *Cth*^−/−^ mice fed CTRL or CysF diet for 6 days (n=4 *Cth*^−/−^ CTRL, n=5 *Cth*^−/−^ CysF). l) Schematic summary of changes in the metabolites in the serum of *Cth*^−/−^ mice fed CTRL or CysF diet for 6 days. Blue lines represent measured, but unchanged metabolites, red and green arrows indicate significantly decreased or increased metabolites, respectively (p<0.05). m) Box plots of GSSG and threonine quantification in the SFAT of *Cth*^−/−^ mice fed CTRL or CysF diet for 6 days (n=6/group). n-o) RNA-seq based expression of (n) *Gclc*, *Gss* and (o) *Bola3* in the SFAT of *Cth*^−/−^ mice fed with CTRL or CysF for 6 days. p) Analysis of EPR spectra of POBN-lipid radical adducts measured in Folch extracts of VFAT, SFAT and BAT tissues from *Cth*^−/−^ mice fed with CTRL or CysF diet for 5 days, normalized to 100 mg (n.d=not detectable, n=5-6/group). q) Aconitase activity determined in VFAT, SFAT and BAT tissues from Cth^−/−^ fed with CTRL or CysF diet for 5 days (n=6-7/group). Data are represented as mean ± SEM. Unless mentioned, differences were determined with unpaired t-tests (*p<0.05, **p<0.01, ***p<0.001).

Metabolomic analyses revealed that despite an increase in *CTH* expression post-CR, adipose cysteine levels were significantly reduced upon CR (Fig. 1f) with no change in homocysteine and cystathionine (Extended Data Fig 1a). Consistent with the reduced expression of *BHMT*, there was a decline in concentration of dimethylglycine (DMG) (Fig. 1f). CR caused a reduction in cysteine derived metabolites, γ-glutamyl-cysteine (γ-Glu-Cys), glutathione (GSH), and cysteinylglycine (Cys-Gly) (Fig. 1g). Collectively, these results suggests that CR in humans reduces enzymes and metabolites that feed into methionine cycle and lowers cysteine (Fig. 1c).

### Cysteine depletion causes lethal weight loss in mice

Cysteine is thought to be biochemically irreplaceable because methionine, the other sole proteinogenic SAA, lacks a thiol group and hence cannot form complexes with metals to control redox chemistry^22^. To determine whether cysteine is required for survival and organismal metabolism, we created a loss of function model where cysteine becomes an essential amino acid requiring acquisition from the diet by deletion of CTH (*Cth^−/−^* mice) (Fig. 1h and Extended data Fig. 1b). Cysteine deficiency was thus induced by feeding adult *Cth^−/−^* mice a custom amino acid diet that only lacks cystine (CysF diet), while control mice were fed an isocaloric diet that contained cystine (CTRL diet) (Fig. 1h). Utilizing this model, we found that mice with cysteine deficiency rapidly lost ∼25-30% body weight within 1 week compared to littermate *Cth*^+/+^ mice fed a CysF diet or *Cth*^−/−^ fed a control diet (Fig. 1i, Extended data Fig. 1b). Upon clinical examination of the cysteine deficient mice, 30% weight loss is considered a moribund state that required euthanasia. The weight loss in mice lacking CTH and cystine in the diet was associated with significant fat mass loss relative to lean mass (Extended data Fig. 1c) in cysteine-deficient animals. Pair feeding of cysteine-replete mice with cysteine depleted diet fed animals produced similar weight-loss (Extended data Fig. 1e). This rapid weight loss is not due to malaise or behavioral alteration, as *Cth*^−/−^CysF mice displayed normal activity and a slight reduction in food intake in the first 2 days after CysF diet switch that was not significantly different (Extended data Fig. 1f and link of video file of cage activity). The *Cth* deficient mice on the control diet were indistinguishable from control littermates in parameters indicative of health, they displayed higher nest building and no change in grip strength, gait, ledge test, hindlimb clasping, and displayed no clinical kyphosis (Extended data Fig. 1g, h). Furthermore, compared to *Cth^−/−^* mice on control diet, the analyses of liver, heart, lungs, and kidneys of *Cth^−/−^* CysF mice did not reveal pathological lesions indicative of tissue dysfunction (Extended data Fig. 1i). Notably, restoration of up to 75% cysteine levels in the diet of *Cth*^−/−^ CysF mice that were undergoing weight-loss was sufficient to completely rescue the body weight over three weight-loss cysteine depletion cycles, demonstrating the specificity and essentiality of cysteine for the organism (Fig. 1j).

To identify systemic changes in metabolites upon cysteine deficiency, we conducted serum and adipose tissue metabolomics analyses. Compared to *Cth-*deficient mice fed a normal diet, the *Cth^−/−^* CysF mice had reduced cystine levels, suggesting that cysteine deficiency is maintained by a reduction in systemic cystine levels (Fig. 1k). Cysteine depletion also elevated the cystathionine and L-serine levels, compared to control diet fed animals (Fig. 1k). Other sulfur amino acid (SAA) metabolites such as methionine, homocysteine (HCys) and glutamic acid were not significantly changed (Extended data Fig. 1j). Taurine levels in the Cth deficient mice on cysteine free diet also did not change compared to control animals (data not shown). Interestingly, the gamma-glutamyl peptide analogs of cysteine and GSH such as 2-aminobutyric acid (2AB) and ophthalmic acid (OA or γglutamyl-2AminobutyrylGlycine) were increased in the serum of cysteine deficient mice (Fig. 1k). Notably, in subcutaneous adipose tissue, cysteine deficiency did not affect glutathione (GSH) (Extended data Fig. 1k) but lowered oxidized GSH (GSSG) concentration, a key downstream product derived from cysteine in TSP (Fig. 1l, m). The increase in γ-glutamyl peptides (2AB and OA) in cysteine-limiting conditions *in vivo* is consistent with studies that show that GCLC can synthesize γglutamyl-2AminobutyrylGlycine in a GSH independent manner and prevents ferroptosis by lowering glutamate generated oxidative stress^23^. OA is a GSH analog in which the cysteine group is replaced by L-2-aminobutyrate (2AB). 2oxobutyrate is the canonical substrate for 2AB in cysteine-replete conditions such that 2AB is produced from 2OB and glutamate in the presence of aminotransferases^24^. Thus, the increase in 2AB despite the removal of cysteine in diet could be due to an alternative pathway of deamination of threonine into 2AB^25^. Indeed, L-threonine levels are increased upon cysteine depletion in mice (Fig. 1m). Prior studies found that GSH can inhibit glutamate cysteine ligase (GCLC)^26,27^ regulating its production by a feedback mechanism. Thus, the removal of cysteine and reduction of GSH may release this disinhibition (Fig 1l). Consistent with this hypothesis and elevated OA levels, *Gclc* and *Gss* expression were increased in cysteine-starved mice (Fig. 1n). The increased OA production vs GSH production reveals adaptive changes induced by systemic cysteine deficiency. Cysteine is also required for Fe-S clusters in numerous proteins^18,28^. We found that cystine-depletion upregulates *Bola3* (Fig. 1o) and *Isca1* gene expression in adipose tissue without affecting *Nfs1* (Extended data Fig. 1l), which are implicated in FeS cluster formation^28^. Consistent with the association between increased *Bola3* and adipose browning in a cysteine-deficient state, adipose-specific deletion of *Bola3* decreases EE and increases adiposity in mice upon aging^29^. The impact of cysteine starvation on Fe-S cluster formation and function requires further studies. The *in vivo* spin trapping and electron paramagnetic resonance (EPR) spectroscopy revealed that cysteine deficiency significantly increased lipid-derived radicals in BAT with undetectable signals in WAT (Fig 1p, Extended Data 1m). Also, given aconitase is regulated by reversible oxidation of (4Fe-4S)^2+^ and cysteine residues, depletion of cysteine also reduced aconitase activity in SFAT with no change in BAT (Fig1q). Together, these data demonstrate that removing cysteine causes lethal weight loss and induces adaptive changes in organismal metabolism, including non-canonical activation of GCLC elevated γ-glutamyl peptides and, GSSG depletion (Fig. 1l).

### Cysteine elimination drives adipose tissue browning

The decrease in fat mass during cysteine deficiency is driven by loss of all major fat depots including subcutaneous fat (SFAT), visceral epididymal/ovarian adipose fat (VFAT), and brown adipose tissue (BAT) (Extended data Fig. 2a). Histological analyses revealed that this reduction in adipose tissue size is associated with transformation of white adipose depots into a BAT-like appearance, with the formation of multilocular adipocytes, enlarged nuclei, and high UCP1 expression, a phenomenon known as ‘browning’ that increases thermogenesis^3,4^ (Fig. 2a, b Extended data Fig. 2b). Interestingly, the SFAT browning in cysteine-deficient mice was reduced upon cysteine-restoration in diet (Fig. 2b). Similar response was observed in visceral fat (VFAT) (Extended data 2b). Consistent with the browning of SFAT, the cysteine-deficient animals show significantly increased expression of UCP1 (Fig 2c) and thermogenic marker genes (Fig. 2d). The UCP1 and ATGL induction upon cysteine-deficiency in adipose tissue was reversed by cysteine-repletion (Fig. 2c). Consistent with 30% weight-loss at day 5, the glycerol concentrations were depleted in the sera of cysteine-deficient mice and were restored by cysteine-repletion induced weight regain (Extended data Fig. 2c). The differentiation of Cth-deficient preadipocytes to mature adipocytes and subsequent exposure to cysteine-free media did not affect thermogenic genes or UCP1, suggesting that a non-cell autonomous mechanism may control adipocyte browning (Extended data Fig. 2d).

**Figure 2:**
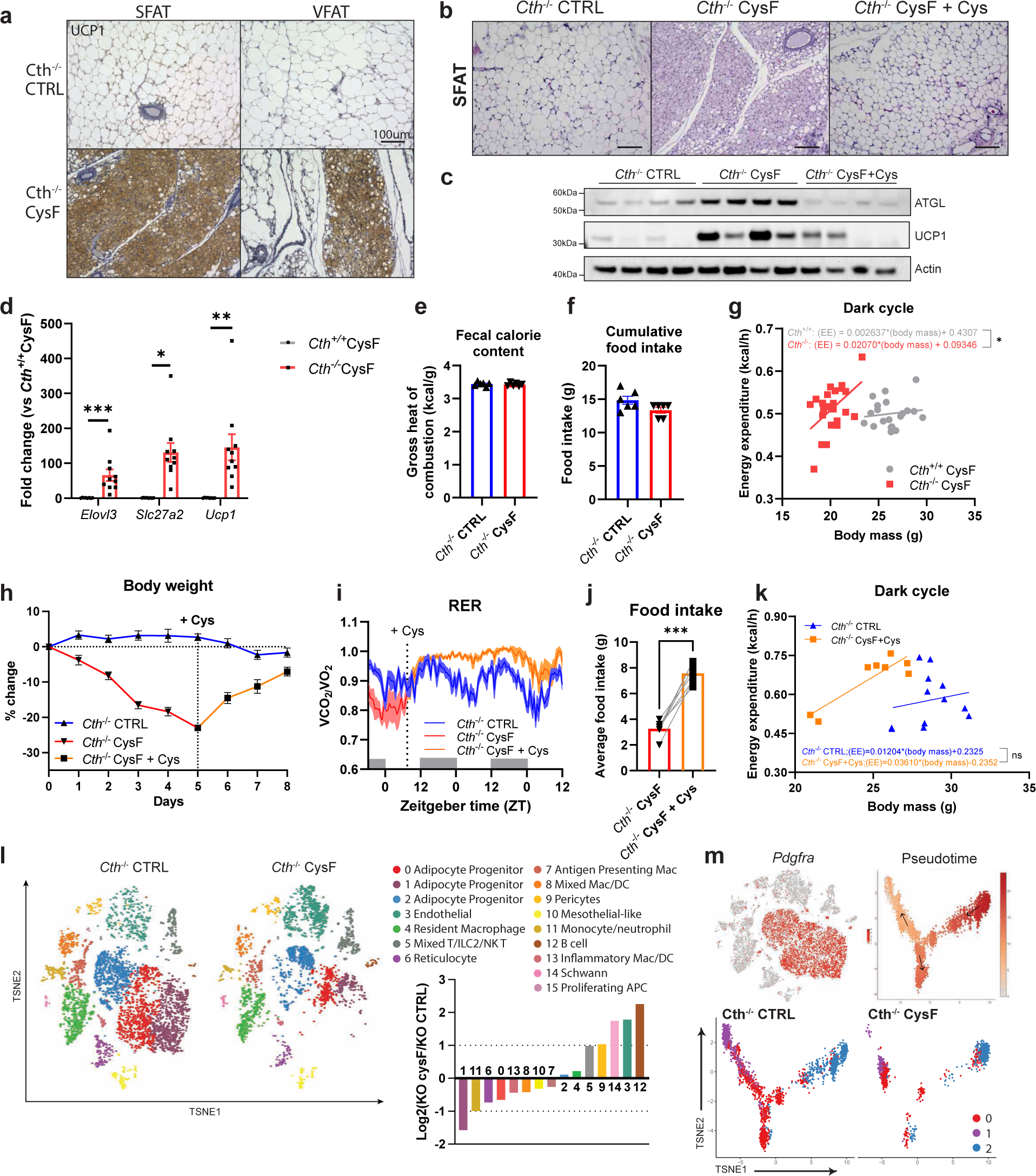
Cysteine depletion induces browning of adipose tissue. a) Representative images of subcutaneous (SFAT) and visceral (VFAT) fat sections stained for UCP1 from *Cth*^−/−^ mice fed CTRL or CysF diet for 6 days (scale bar=100um). b) Representative H&E-stained sections of SFAT of *Cth*^−/−^ mice fed CTRL or CysF diet for 6 days or CysF diet followed by Cys-supplemented diet for 4 days (CysF+Cys) (scale bar=100 μm). c) Western blot detection of ATGL and UCP1 in SFAT from *Cth*^−/−^ mice after 6 days of CTRL or CysF diet or Cys supplementation after CysF-induced weight loss. Actin is used as a loading control. d) qPCR analysis of thermogenic genes in SFAT of *Cth*^+/+^ and *Cth*^−/−^ mice fed CysF diet for 6 days (n=8 *Cth^+/+^* and n=10 *Cth^−/−^*). e) Fecal calorie content and f) cumulative food intake of *Cth*^−/−^ mice fed CTRL or CysF diet for 4 days (n=6/group). g) Linear regression analysis of energy expenditure against body mass during dark cycle at 4, 5 days of weight loss (n=10 *Cth*^+/+^ CysF and n=12 *Cth*^−/−^ CysF). h) Percent body weight change of *Cth*^−/−^ mice fed with CTRL diet or CysF diet (red line) for 5 days and then switched to Cys-containing diet (orange line) for 3 days (n=6/group). i) Respiratory exchange ratio (RER) measured in metabolic cages, of *Cth*^−/−^ mice fed with CTRL diet or Cys-containing diet after CysF induced weight loss (n=4-6/group). j) Average food intake of *Cth*^−/−^ mice fed with CysF diet and then switched to Cys-containing diet for 2 days (n=7/group). Significance was measured with paired t-test. k) Linear regression analysis of energy expenditure against body mass during dark cycle of *Cth*^−/−^ mice fed with CTRL or Cys-supplemented diet after CysF induced weight loss (n=4-6/group), average values of the first two nights after diet switch. l) t-SNE plot of scRNAseq showing cluster identities from SFAT stromal vascular fraction from *Cth*^−/−^ mice fed CTRL or CysF diet at day 4 of weight-loss and bar chart showing population fold changes in relative abundance of each cluster comparing *Cth*^−/−^ CysF vs. *Cth*^−/−^ CTRL. m) t-SNE plot displaying *Pdgfra* expression in red across all populations and monocle analysis of clusters 0, 1, and 2, with coloring by pseudotime to show right most cluster giving rise to two separate clusters. Each cluster represented by color in *Cth*^−/−^ CTRL and *Cth*^−/−^ CysF. Data are expressed as mean±SEM. Statistical differences were calculated by 2-way ANOVA with Sidak’s correction for multiple comparisons or unpaired t-test (*p<0.05, **p<0.01, ***p<0.001).

We next investigate whether energy absorption, energy-intake or energy expenditure contributes to the cysteine-depletion induced weight-loss. Analysis of energy absorption by fecal bomb calorimetry revealed no significant difference in control and cysteine-deficient mice (Fig. 2e). Moreover, although the cumulative food intake over 5 days of weight loss was not statistically different, the cumulative food intake in the first 2 days (Extended data Fig. 2d) after switching to CysF diet was lower (p < 0.05) which may contribute to early weight loss. Calculation of the analysis of covariance (ANCOVA) or representation of the data as regression between energy expenditure and body mass^5,7^, demonstrated that EE is increased in cysteine deficient animals during the dark cycle (Fig. 2g) and not in the light cycle (Extended data Fig. 2f, g). In addition, there was no difference in locomotor activity between control or cysteine-deficient mice (Extended data Fig. 2h), suggesting cysteine depletion increases EE. Moreover, the increase in EE was supported by increased fat utilization, as the respiratory exchange ratio (RER) in cysteine-deficient animals was significantly reduced (Extended data Fig. 2i, j).

We next determined the specificity of cysteine on mechanisms that may contribute to rapid weight loss. Interestingly, weight-regain post cysteine repletion significantly reversed adipose-browning (Fig 2b, Extended data Fig. 2b) and normalized the glycerol, ATGL and UCP1 levels in adipose tissue. (Fig. 2c). Furthermore, cysteine replacement also reversed the cysteine-deficiency-induced reduction in RER, suggesting the restoration of organismal metabolism to carbohydrate utilization instead of fatty acid oxidation (Fig. 2h, i). Surprisingly, cysteine repletion significantly increased food intake for the first two days, suggesting that animals sense cysteine in diet and compensate via hyperphagia to restore bodyweight setpoint (Fig. 2j). The EE upon cysteine-replacement was not significantly different during weight rebound (Fig. 2k). These data suggest that cysteine replacement can rapidly reverse weight loss by mechanisms that involve reduced adipose browning, decreased fat utilization as well as increased energy intake.

We conducted the RNA-sequencing of the major adipose depots to investigate the mechanisms that control adipose tissue browning and associated remodeling. As displayed by the heatmap, cysteine deficiency profoundly alters the transcriptome of adipose tissue (Extended data Fig. 2k). Gene set enrichment analysis comparing *Cth*^−/−^ CTRL vs *Cth^−/−^* CysF identified that the top downregulated pathways are involved in the extracellular matrix and collagen deposition, highlighting the broad remodeling of the adipose tissue (Extended data Fig. 2l). In addition, multiple metabolic pathways appear to be regulated by cysteine deficiency within the SFAT with ‘respiratory electron transport chain and heat production’ as the top pathway induced during cysteine deficiency (Extended data Fig. 2l). Indeed, numerous genes identified by the ‘thermogenesis’ GO-term pathway such as *Ucp1, Cidea, Cox7a1, Cox8b, Dio2, Eva1, Pgc1, Elovl3,* and *Slc27a2,* are differentially expressed comparing *Cth*^+/+^ CysF and *Cth*^−/−^ CysF in the SFAT (Extended data Fig. 2m). These results demonstrate that cysteine depletion activates the thermogenic transcriptional program.

To investigate the cellular basis of adipose tissue remodeling during cysteine deficiency, we isolated stromal vascular fraction (SVF) by enzymatic digestion and conducted single-cell RNA sequencing of SFAT. We isolated SVF cells from *Cth*^+/+^ and *Cth*^−/−^ fed CTRL or CysF diet with each sample pooled from 4 animals (Extended data Fig. 3a). A total of 4,666 cells in *Cth*^+/+^ CTRL; 5,658 cells in *Cth*^+/+^ CysF; 4,756 cells in *Cth*^−/−^ CTRL; and 3,786 cells in *Cth*^−/−^ CysF were analyzed for scRNA-seq (Extended data Fig. 3b). Consistent with prior results^30,31^, the unbiased clustering revealed 15 distinct cell populations including αβ T cells, γδ T cells, ILC2s, and NK T cells, B cells, reticulocytes, mesothelial-like cells, Schwann cells, and several myeloid clusters (Extended data Fig. 3b-d). Comparison of *Cth*^−/−^ CysF with other groups revealed dramatic changes in cellular composition (Fig. 2l). Particularly, loss of clusters 0, 1, and 2 were apparent upon cysteine deficiency (Fig. 2l). Furthermore, these clusters contained the highest numbers of differentially expressed genes induced by β3-adrenergic receptor agonist CL-316243^32^ (Extended data Fig. 3e), highlighting them as important cell populations in regulating the effects of cysteine deficiency. By expression of *Pdgfra*, we identified these clusters as adipocyte progenitors (Fig. 2h). We conducted a pseudo-time analysis to place these clusters on a trajectory and illuminate their cell lineage. Trajectory analysis based on pseudo-time suggested that cluster 2 may differentiate into two separate preadipocyte clusters, clusters 0 and 1 (Fig. 2m). *Cth*^−/−^ CysF animals proportionally lost Clusters 0 and 1, while relatively maintaining cluster 2 compared to the other groups (Fig. 2m), suggesting that more differentiated preadipocytes are mobilized during cysteine deficiency. Indeed, cluster 2 expressed *Dpp4*, an early progenitor marker that has been shown to give rise to different committed preadipoctyes^33^ (Extended data Fig. 3f). Cluster 0 was enriched for both *Icam1* and *F3*, which are expressed by committed adipogenic, and antiadipogenic preadipocytes, respectively ^30,33^ (Extended data Fig. 3g, h). *Cd9*, a fibrogenic marker in preadipocytes ^32,34^, along with the collagen gene, *Col5a3,* were broadly expressed across clusters 0 and 1, and was specifically lost by day 4 of inducing cysteine deficiency (Extended data Fig. 3g). The loss of these preadipocyte clusters were orthogonally validated by FACS (Extended data Fig. 3h). We next sought to identify beige/brown adipocyte precursors in our scRNA-seq dataset to understand whether there was an increased commitment towards brown adipocytes. Clearly, *Tagln,* or Sm22, which has been previously described in beige adipocytes^35,36^, is specifically expressed by a subset of cells in cluster 1 (Extended data Fig. 3g). Interestingly, these *Tagln*-expressing cells are lost with cysteine deficiency (Fig. 2i). Given the strong browning phenotype observed on day 6, it is possible that these cells become mobilized and differentiate early on during cysteine deficiency, leading to the absence of these cells as mature adipocytes are not captured within the SVF. Indeed, when we performed pathways analysis on cluster1, comparing gene expression of *Cth*^−/−^ CysF with *Cth*^−/−^ CTRL, we found that one of the top upregulated pathways was ‘adipogenesis’ (Extended data Fig. 3i). Furthermore, examination of the expression of stem associated markers and mature adipocyte markers in the adipocyte progenitor clusters revealed a clear downregulation of stem markers and an increase in mature adipocyte markers, suggesting that cysteine deficiency was driving the maturation of progenitor cells (Fig. 2m and Extended data Fig. 3j). However, given the robust transformation of the adipose tissue during cysteine deficiency towards browning, it is unlikely that mobilization of brown precursors alone is mediating this response. Prior studies have found that in certain models, beige adipocytes can originate from pre-existing white adipocytes, in addition to de-novo adipogenesis^37^. The potential role of cysteine in the trans-differentiation of mature white adipocytes into brown-like adipocytes needs to be further examined using future lineage-tracking studies.

**Figure 3:**
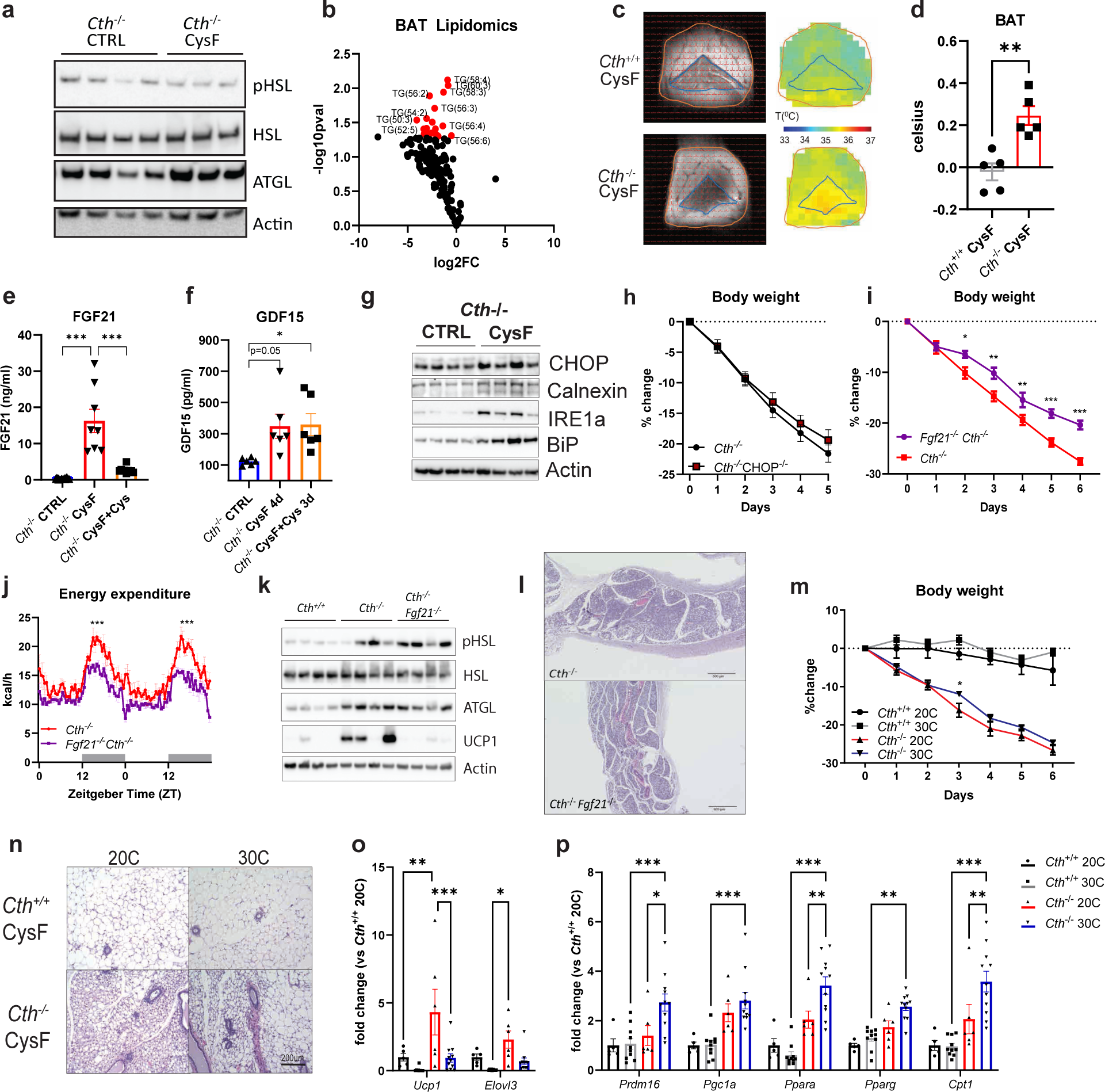
FGF21 is partially required for cysteine-restriction mediated weight-loss. a) Western blot detection of lipolysis regulators pHSL, HSL and ATGL in SFAT from *Cth*^−/−^ mice after 6 days of CTRL or CysF diet, actin is used as loading control. b) Volcano plot of lipid species of BAT showing fold change of triglycerides in *Cth*^−/−^ mice fed CTRL or CysF diet. c) *in vivo* measurement of BAT temperature by BIRDS imaging and d) quantification of local temperature differences in BAT compared to surrounding tissue in *Cth*^+/+^ and *Cth*^−/−^ mice on CysF diet for 6 days (n=5/group). e) Serum FGF21 quantification in *Cth*^−/−^ CTRL (n=23), *Cth*^−/−^ CysF for 6 days (n=8) and Cth^−/−^ CysF followed with 4 days of Cys supplementation (n=10). f) Serum GDF15 concentrations in *Cth*^−/−^ CTRL, *Cth*^−/−^ CysF for 4 days and Cth^−/−^ CysF followed with 3 days of Cys supplementation (n=6/group). g) Immunoblot analysis of CHOP, Calnexin, IRE1a, BiP in the liver of *Cth*^−/−^ mice fed with CTRL or CysF diet at day 6. Actin was used as loading control. h) Percentage body weight change of *Cth*^−/−^ and *Cth*^−/−^CHOP^−/−^ mice fed with CysF diet for 5 days (n=17 *Cth*^−/−^ and n=15 *Cth*^−/−^CHOP^−/−^). i) Percentage body weight change of *Cth*^−/−^ and *Fgf21*^−/−^*Cth*^−/−^ mice fed with CysF diet for 5 days (n=13 *Cth*^−/−^ and n=18 *Fgf21*^−/−^*Cth*^−/−^). j) Energy expenditure measured in metabolic cages of *Cth*^−/−^ and *Cth*^−/−^ *Fgf21*^−/−^ mice on days 3-4 of CysF diet (n=5/group). k) Immunoblot analysis of pHSL, HSL, ATGL, and UCP1 in SFAT of *Cth*^+/+^, *Cth*^−/−^ and *Cth*^−/−^*Fgf21*^−/−^ mice fed CysF diet for 6 days. l) Representative H&E stained SFAT sections of *Cth*^−/−^ and *Fgf21*^−/−^*Cth*^−/−^ mice after 6 days of CysF diet (scale bar=500um). m-p) *Cth*^+/+^ and *Cth*^−/−^ mice were fed with CysF diet and housed at 20°C or 30°C for 6 days. m) Percentage body weight change (n=3 *Cth*^+/+^ 20°C, n=4 *Cth*^+/+^ 30°C, n=4 *Cth*^−/−^ 20°C, n=5 *Cth*^−/−^ 30°C), n) representative images of H&E staining of SFAT sections (scale bar=200um) and o-p) qPCR analysis of thermogenic markers (n=5 *Cth*^+/+^ 20°C, n=10 *Cth*^+/+^ 30°C, n=6 *Cth*^−/−^ 20°C, n=11 *Cth*^−/−^ 30°C). Data are expressed as mean±SEM. Statistical differences were calculated by one-way ANOVA with Tukey’s correction for multiple comparisons or 2-way ANOVA with Sidak’s correction for multiple comparisons or unpaired t-test (*p<0.05, **p<0.01, ***p<0.001).

### Cysteine depletion-induced FGF21 is partially required for weight loss

To determine the mechanism of adipose thermogenesis caused by cysteine starvation, we next investigated the processes upstream of increased fatty acid oxidation. We measured the lipolysis regulators pHSL and ATGL and found that cysteine deficiency increases ATGL expression without consistently affecting pHSL levels (Fig. 3a, Extended data Fig. 4a). ATGL preferentially catalyzes the first step of triglyceride hydrolysis whereas HSL has a much broader range of substrates with a preference for diacylglycerols and cholesteryl esters^38^. Given a dramatic browning response in WAT post-cysteine deficiency, the increased ATGL is consistent with prior work that shows BAT relies heavily on the action of ATGL to mobilize lipid substrates for thermogenesis^39^. This is further supported by a decrease in most lipid species, particularly triglycerides and diacylglycerol in the BAT of cysteine deficient mice (Fig. 3b, Extended data Fig. 4b, c). Considering dramatic adipose tissue browning and elevated UCP1 expression upon cysteine starvation, we next sought to investigate whether this is a homeostatic response to defend core-body temperature (CBT) or if temperature set-point is perturbed to causes hyperthermia. We measured core body temperature utilizing loggers surgically implanted into the peritoneal cavity in *Cth*^−/−^ mice on CTRL or CysF diet over 6 days period when animals lose weight. Surprisingly, despite conversion of WAT into brown-like thermogenic fat, the core body temperature was not different between control and cysteine deficient mice (Extended data Fig. 4d, e). These data suggest that either cysteine-may signal the host to defend CBT within tight normal physiological range or any metabolic heat that is generated is dissipated due to the animal housing in the subthermoneutral temperature. To further confirm adipose thermogenesis *in vivo*, we utilized a highly sensitive and specific magnetic resonance spectroscopic imaging (MRSI) method called Biosensor Imaging of Redundant Deviation in Shifts (BIRDS)^40^ to determine the temperature of BAT in *Cth*^+/+^ and *Cth*^−/−^ animals after 6 days of CysF diet. This method relies on measuring the chemical shift of the four non-exchangeable methyl groups from an exogenous contrast agent, TmDOTMA, which has a high-temperature sensitivity (0.7 ppm/°C). The TmDOTMA^−^ methyl resonance has ultra-fast relaxation times (<5ms), allowing high signal-to-noise ratio by rapid repetition for superior signal averaging^40^. The temperature was calculated from the chemical shift of the TmDOTMA^−^ methyl resonance according to (eq. 1 methods). Compared to cysteine-replete animals, the *in vivo* local temperature in BAT of cysteine-deficient mice was significantly greater than surrounding tissue (Fig. 3c, d), suggesting increased thermogenesis.

**Figure 4:**
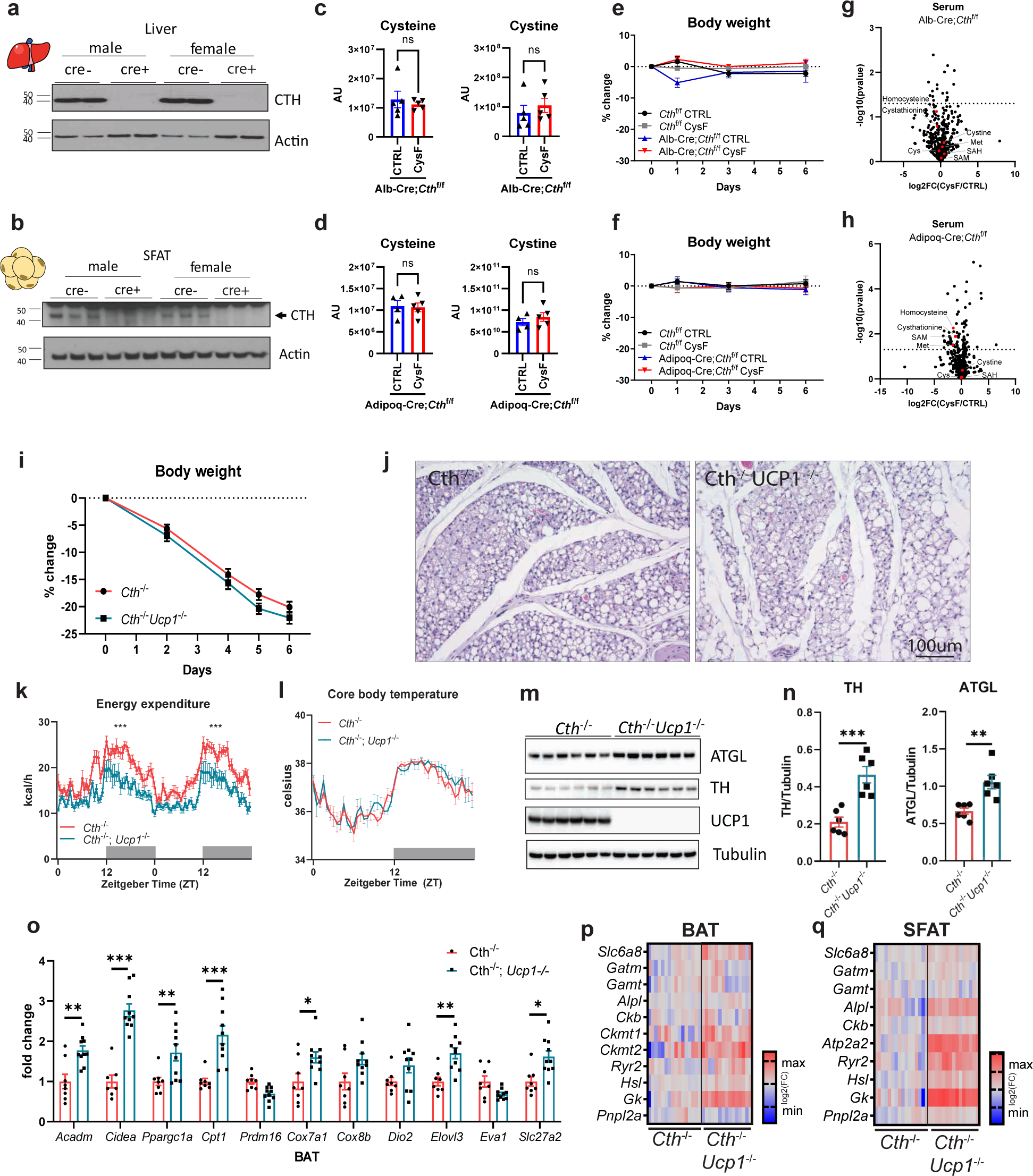
Global cysteine deficiency induced adipose browning is UCP1 independent. a) Immunoblot analyses of CTH in the liver of male and female *Cth*^f/f^ Alb:Cre^−^ or Alb: Cre^+^ mice. d) Western blot detection of CTH in the SFAT of male and female *Cth*^f/f^ Adipoq:Cre^−^ or Adipoq:Cre^+^ mice. c-d) Serum cysteine and cystine determined by LC-MS/MS in c) Alb:Cre^+^*Cth*^f/f^ mice and d) Adipoq:Cre;*Cth*^f/f^ mice after 5 days of CTRL or CysF diet (n=4-5/group). AU: arbitrary units. e-f) Percentage body weight changes of e) Alb-Cre;*Cth*^f/f^ mice and f) Adipoq-Cre;*Cth*^f/f^ mice after 5 days of CTRL or CysF diet (n=4-5/group). g-h) Volcano plot of serum metabolites identified by LC-MS/MS in g) Alb-Cre;*Cth*^f/f^ mice and i) Adipoq-Cre;*Cth*^f/f^ mice after 5 days of CTRL or CysF diet (n=4-5/group). Transsulfuration pathway related metabolites are highlighted in red. Cys: cysteine. Met: methionine. SAH: S-adenosyl homocysteine. SAM: S-adenosyl methionine. i-k) *Cth*^−/−^ and *Cth*^−/−^ *Ucp*1^−/−^ mice were fed a CysF diet for 6 days (n=8/group). i) Percent body weight change over 6 days of diet. j) Representative H&E histology images of SFAT after 6 days of diet. k) Energy expenditure measured in metabolic cages on days 4 and 5 of CysF diet. l) Core body temperatures (CBT) measured in the peritoneal cavity by implantation of Star-Oddi loggers over 6 days of diet in male *Cth*^−/−^ and *Cth*^−/−^ *Ucp1*^−/−^ mice fed CysF diet. Recordings were taken every 30min and representative day 4 is plotted (n=7 *Cth*^−/−^, n=5 *Cth*^−/−^ *Ucp1*^−/−^). m) Immunoblot staining of ATGL, TH, and UCP1 in BAT of *Cth*^−/−^ and *Cth*^−/−^ *Ucp1*^−/−^ fed a CysF diet for 6 days and n) quantification using tubulin as loading control. o) Thermogenic markers gene expression analysis in BAT of *Cth*^−/−^ and *Cth*^−/−^ *Ucp1*^−/−^ mice fed a CysF diet for 6 days, measured by qPCR (n=8 *Cth*^−/−^, n=10 *Cth*^−/−^ *Ucp1*^−/−^). p-q) Heatmaps of gene expression of genes involved in creatine, calcium and lipid futile cycles in p) BAT and q) SFAT of *Cth*^−/−^ and *Cth*^−/−^ *Ucp1*^−/−^ mice fed a CysF diet for 6 days (n=15-16/group), quantified by qPCR. Data are expressed as mean±SEM. Statistical differences were calculated by 2-way ANOVA with Sidak’s correction for multiple comparisons, or by unpaired t-test (*p<0.05, **p<0.01, ***p<0.001).

Changes in nutritional stress induced by caloric restriction, methionine restriction, or low protein diets upregulate the expression of FGF21, which, when overexpressed, increases lifespan and also upregulates EE^41,42^. The induction of cysteine deficiency in *Cth* deficient mice caused a dramatic increase in the FGF21 concentration in blood (Fig. 3e) and *Fgf21* expression in the liver (Extended data Fig. 4f), which was reversed by cysteine-repletion induced weight regain (Fig 3e). Similar to FGF21, the hormone GDF15, can also be induced by cellular or nutritional stress-mediated signaling^43^. Cysteine depletion at day 4 post-weight loss significantly increased GDF15, which was not restored after cysteine-repletion-induced weight regain (Fig 3f). Future studies are required to determine if GDF15 is dispensable for cysteine-depletion-induced weight loss. Given the cysteine-repleted diet switch increases food intake, the higher GDF15 levels during weight-rebound are likely insufficient to cause food aversion. Recent studies suggest elevated endoplasmic-reticulum (ER) stress in *Bhmt^−/−^* mice with reduced methionine cycle, is associated with increased FGF21 and adipose browning^44^. Notably, cysteine deficiency led to induction of ER stress proteins CHOP, Calnexin, IRE1α and BIP (Fig. 3g). However, deletion of CHOP in cysteine-starved mice did not rescue weight-loss (Fig. 3h) or affected the FGF21 and GDF15 serum levels (Extended data Fig. 4g,h) suggesting that CHOP dependent ER-stress response does not drive cysteine’s neuroendocrine or metabolic effect. Given cysteine specifically regulated FGF21 during weight loss and regain (Fig 3e), we generated *Fgf21^−/−^ Cth^−/−^* DKO mice. In the absence of FGF21, cysteine deficiency-induced weight-loss and reduction in adiposity in *Cth^−/−^* mice were blunted, but the weight-loss trajectory continued and was not rescued (Fig. 3i, Extended data Fig. 4i). The *Fgf21^−/−^ Cth^−/−^* DKO mice had lower EE compared to *Cth^−/−^* mice on CysF diet (Fig. 3j). However, RER was not different, indicating that *Fgf21^−/−^ Cth^−/−^* mice still significantly utilized fat as an energy substrate (Extended data Fig. 4j). This was supported by maintenance of lipolysis signaling observed by levels of pHSL and ATGL in *Cth*^−/−^ mice, but reduced UCP1 protein and mRNA expression in WAT of *Fgf21^−/−^ Cth^−/−^* (Fig. 3k, Extended data Fig. 4k). Surprisingly, the WAT of *Fgf21^−/−^ Cth^−/−^* DKO mice maintained classical multilocular browning characteristics (Fig. 3l) suggesting that FGF21 is not required for adipose browning. These results suggest that FGF21 is partially required for weight loss but does not mediate lipid mobilization or adipose browning caused by cysteine deficiency.

### Cysteine-starvation-induced weight-loss is maintained at thermoneutrality

Cysteine elimination revealed a metabolic crisis that may signal the host to activate thermogenic mechanisms. However, across animal vivaria, including ours, mice are housed at sub-thermoneutral 20°C temperatures and are constantly under thermogenic stress due to slight cold challenge^4^. To further confirm that mice were indeed inducing thermogenesis to defend core body temperature, we housed cysteine deficient animals at 30°C thermoneutrality. The cysteine deficiency in *Cth*^−/−^ mice housed at 30°C also led to similar weight loss as 20°C with significant browning of adipose tissue (Fig. 3m, n, Extended data Fig. 4l). The degree of browning and gene expression of *Ucp1* and *Elovl3* in CysF *Cth* deficient mice at thermoneutrality was relatively lower than inductions observed at 20°C (Fig. 3o). Furthermore, expression of genes involved with lipid regulation and browning such as *Prdm16, Ppargc1a, Ppara, Pparg,* and *Cpt1* (Fig. 3p) were significantly increased in SFAT, suggesting that even at thermoneutral temperatures, *Cth*^−/−^ CysF fed mice activate fat metabolism and, have increased thermogenesis caused by cysteine deficiency. In addition, compared to controls, the cysteine deficient mice at thermoneutrality retained higher UCP1 expression in BAT (Extended data Fig. 4m). Together, cysteine-depletion induced weight loss and adipose browning are maintained at thermoneutrality.

### Systemic depletion of cysteine drives browning in a UCP1-independent manner

The liver is believed to be the primary organ for the maintenance of organismal cysteine homeostasis^6,16^. Immunoblot analyses revealed the highest CTH expression in the liver followed by the kidney, thymus, and adipose tissue (Extended data Fig. 5a). Given CR in humans lowers cysteine in adipose tissue; we next generated adipocyte as well as hepatocyte-specific Cth deficient mice to determine cell type-specific mechanism of cysteine in weight-loss (Fig. 4 a-f). As expected, deletion of *Cth* in the liver did not affect the expression in the kidney, and adipose-specific ablation of *Cth* maintained the expression in the liver (Extended data Fig. 5b). Neither liver nor adipose-specific deletion of *Cth* caused a reduction in serum cysteine levels (Fig. 4c, d and Extended data Fig. 5c,d) or fat-mass loss when cysteine was restricted in the diet (Fig. 4e, f). The further LC/MS analyses of sera of hepatocyte-specific Cth deficient mice maintained on CysF diet had no change in cystathionine, γ-glutamyl-dipeptides, cysteine or cystine (Fig 4 g, Extended data Fig 5e). Consistent with low CTH activity, livers of the CysF-fed mice (AlbCre:*Cth*^f/f^, CysF) had lower levels of cysteine, cystathionine, s-adenosyl homocysteine, 2AB and ophthalmate (Extended data Fig. 5f, g). Furthermore, cystathionine and cysteine/cystine in subcutaneous adipose tissue of liver-specific Cth deficient mice were unchanged (Extended data Fig 5h, i) suggesting specificity of TSP response in liver. Consistent with these data, no change in serum cysteine/cystine were detected in adipose tissue specific Cth^−/−^ mice maintained on cysteine free diet (Fig. 4h, Extended data Fig. 5j). The TSP metabolites can potentially be generated by the gut microbiota^22^. The *Cth*^−/−^ animals co-housed together with *Cth*^+/+^ mice still maintained weight loss when fed a CysF diet, suggesting that microbiota derived metabolites do not account for the weight-loss (Extended data Fig. 5k). These results demonstrates that *Cth* across multiple tissues may defend systemic cysteine concentration to prevent uncontrolled thermogenesis and death when cysteine content is low in diet.

**Figure 5:**
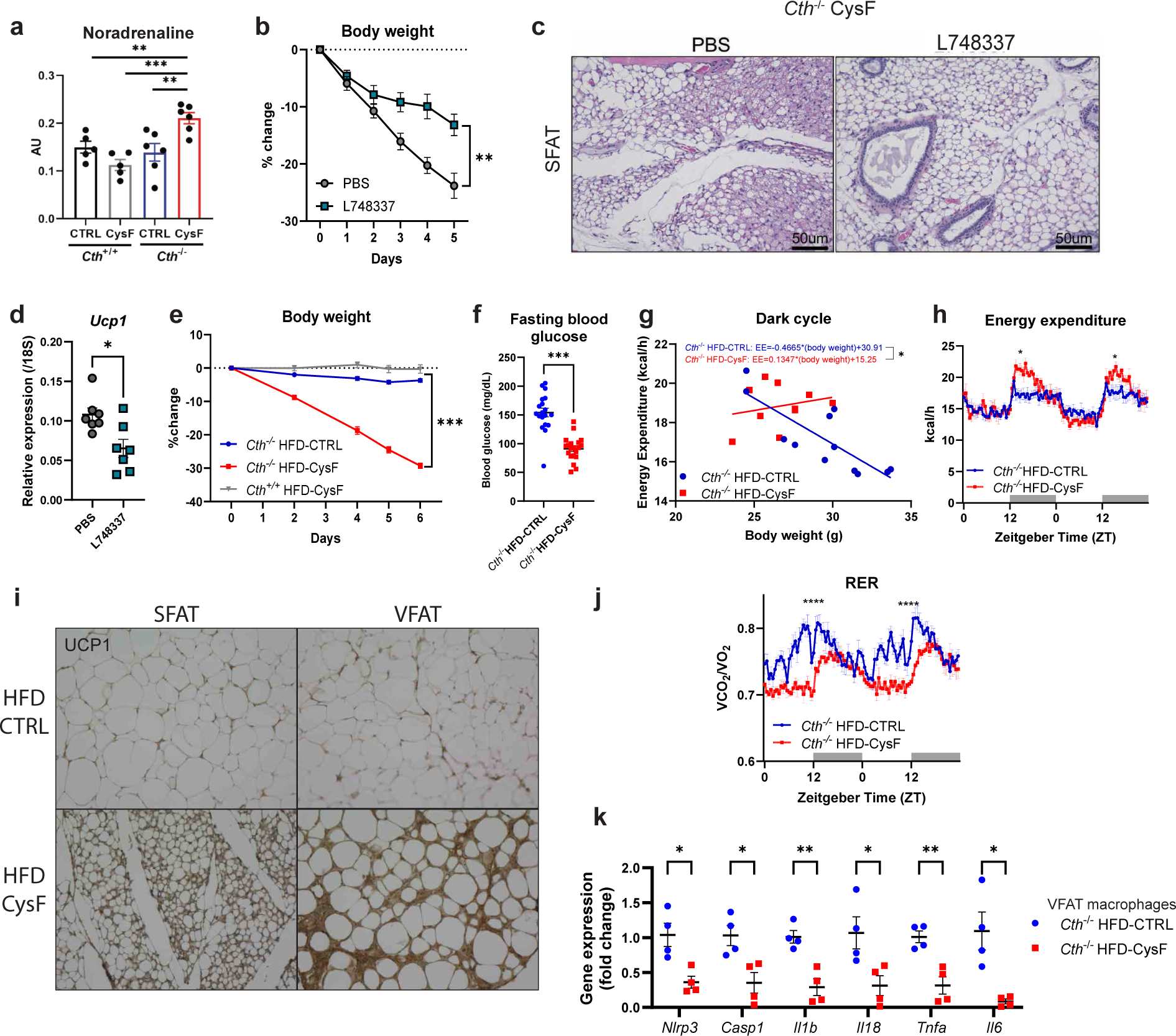
Cysteine-elimination induced browning and weight loss requires noradrenergic signaling. a) Measurement of noradreanaline by orbitrap MS/MS in the SFAT of *Cth*^+/+^ and *Cth*^−/−^ fed 6 days of CTRL or CysF diet (n=5 *Cth*^+/+^ CTRL, n=5 *Cth*^+/+^ CysF, n=6 *Cth*^−/−^ CTRL, n=6 *Cth*^−/−^ CysF). b-d) *Cth*^−/−^ mice were fed with CysF diet for 5 days and treated daily with a β-3 adrenergic receptor antagonist (L748337) or vehicle (PBS) (n=7/group). b) Percentage body weight change. c) Representative images of hematoxylin and eosin (H&E) staining of SFAT sections (scale bar=50um). d) qPCR gene expression of *Ucp1* in BAT depots. e-j) *Cth*^−/−^ mice that had been fed a high fat diet (HFD) for 12 weeks were switched to a HFD containing (HFD-CTRL) or lacking cystine (HFD-CysF). e) Percentage body weight change after switching to HFD-CysF diet (n=6 *Cth*^−/−^ HFD-CTRL, n=5 *Cth*^−/−^ HFD-CysF and n=5 *Cth*^+/+^ HFD-CysF). f) Fasting blood glucose measured 1 week post diet switch (*Cth*^−/−^ HFD-CTRL n=19, *Cth*^−/−^ HFD-CysF, n=20). g) Linear regression analysis of energy expenditure (EE) against body mass during dark cycle and (h) EE of *Cth*^−/−^ mice fed with HFD-CTRL or HFD-CysF, average values of nights 4 and 5 of diet switch (n=6 *Cth*^−/−^ HFD-CTRL, n=5 *Cth*^−/−^ HFD-CysF). i) Representative histological sections of SFAT and VFAT stained for UCP1, 6 days after diet switch. j) Respiratory exchange ratio (RER) measured in metabolic chambers on days 4 and 5 of diet switch (n=6 *Cth*^−/−^ HFD-CTRL, n=5 *Cth*^−/−^ HFD-CysF). k) Q-PCR analysis of inflammatory genes in VFAT macrophages of *Cth*^−/−^ mice after diet switch to HFD-CTRL or HFD-CysF (n=4/group). Data are expressed as mean±SEM. Statistical differences were calculated by 2-way ANOVA with Sidak’s correction for multiple comparisons, or by unpaired t-test (*p<0.05, **p<0.01, ***p<0.001).

Given that UCP1 is a canonical regulator of non-shivering adipose thermogenesis^44,45^ and since cysteine-elimination induced UCP1 expression in WAT, we next deleted UCP1 in cysteine deficient mice to determine its role in adipose browning. Interestingly, we found that *Cth^−/−^Ucp1^−/−^* double knockout (DKO) mice had equivalent food intake (Extended data Fig. 5l) and lost weight at a similar rate to its *Cth^−/−^* littermates on a CysF diet and displayed similar browning-like features with multilocular adipocytes (Fig. 4i, j). The ablation of UCP1 in cysteine-deficient mice lowered EE but did not affect the CBT (Fig. 4k, i). The lack of UCP1 in *Cth* deficient mice undergoing cysteine starvation displayed elevated ATGL and tyrosine hydroxylase (TH) expression, suggesting increased lipolytic signaling (Fig. 4m, n). Despite lack of UCP1, gene expression indicative of the thermogenic program, such as *Ppargc1, Cidea, Cpt1* are significantly increased in *Cth^−/−^ Ucp1^−/−^* DKO mice compared to *Cth^−/−^* in the BAT after 6 days of CysF diet (Fig. 4o). Furthermore, gene expression of other mediators of the thermogenic genes such as *Acadm, Cox7a1, Elovl3*, and *Slc27a2* are also significantly increased in *Cth^−/−^Ucp1^−/−^* DKO mice compared to *Cth*^−/−^ animals fed cysteine-restricted diet (Fig. 4o). The UCP1-independent thermogenesis has been reported^47^. The creatine futile cycling is proposed to regulate UCP1-independent thermogenesis^48^. Compared to control animals, the creatine cycle genes *Ckb* and *Alpl* were not significantly different in SFAT of cysteine-deficient animals (Extended data Fig. 5m). The creatine synthesis genes, *Gatm* and *Gamt* were significantly reduced with cysteine deficiency in the SFAT (Extended data Fig. 5m). The expression of one of the creatine kinases that utilize ATP, *Ckmt2*, and the transporter for creatine, *Slc6a8* were also not differentially regulated in SFAT (Extended data Fig. 5m). Interestingly, *Ckmt1* and *Ckmt2* expression was increased in BAT of *Cth*^−/−^*Ucp1*^−/−^ animals compared to cysteine-deficient animals (Fig. 4p). In addition, alternative UCP1-independent thermogenic regulatory genes *Atp2a2* and *Ryr2* that control calcium cycling^49^ were not impacted by cysteine deficiency (Extended data Fig. 5n). Similarly, *Sarcolipin* and *Atp2a2* that can increase muscle driven thermogenesis^50^ were also not affected in skeletal muscle of *Cth* deficient mice lacking cysteine (Extended data Fig. 5o). Futile lipid cycle is also implicated in UCP1 independent thermogenesis^51^. Interestingly, *Cth*^−/−^ mice on CysF diet have significantly elevated expression of *Dgat1*, *Pnpla2* and *Gk* with no change in *Lipe* in SFAT (Extended data Fig. 5p). The expression of these genes is also induced in absence of UCP1 in SFAT (Fig. 4q). However, absence of association between changes in gene expression of major UCP1 independent regulators does not rule out causal role of some of these mechanisms in cysteine-elimination driven adipose browning. These results suggest that systemic cysteine deficiency-induced thermogenesis depends on a non-canonical UCP1-independent thermogenic mechanism.

### Cysteine depletion-induced adipose browning and weight loss requires catecholamine signaling

Since cysteine-elimination-induced adipocyte browning is non-cell autonomous (Extended data Fig. 2d), we investigated the mechanism of adipose browning.

Upstream of lipolysis, non-shivering thermogenesis is mainly activated by the sympathetic nervous system (SNS) derived adipose noradrenaline^52^. Mass-spectrometric analyses of subcutaneous adipose tissue (Fig. 5a), including imaging mass spectrometry of BAT (Extended data Fig. 6a) revealed that cysteine-starvation induced browning is associated with increased noradrenaline (NA) concentrations. This was coupled with a significant reduction in NA-degrading enzyme monoamine oxidase-a (*Maoa*), without affecting catechol-o-methyl transferase (*Comt*), suggesting increased adipose NA bioavailability (Extended data Fig. 6b,c). Finally, to test whether SNS derived NA is required for adipose browning, the inhibition of β3-adrenergic receptors (ADRB3) by L748337 in *Cth* deficient mice lacking cysteine-protected animals against weight loss (Fig. 5b), blunted adipose browning (Fig. 5c) and lowered browning marker *Ucp1* (Fig. 5d). This was consistent with our unbiased RNA sequencing analyses that showed that cysteine-regulated adipose clusters contained the highest numbers of differentially expressed genes induced by β3-adrenergic receptor agonist (Extended data Fig. 3e). Together our findings suggest that cysteine-depletion drives increased sympathetic activity leading to augmented ADRB3-mediated NA signaling that controls adipose browning to weight loss.

### Cysteine deficiency reverses high-fat diet-induced obesity in mice

We next tested whether cysteine deficiency could be utilized to induce an adaptive thermogenic mechanism for fat mass reduction in the high-fat diet (HFD) induced obesity model. The *Cth^−/−^* mice that had been fed HFD for 12 weeks were switched to an isocaloric HFD containing (HFD-CTRL) or lacking cystine (HFD-CysF). The *Cth^−/−^* mice fed HFD-CysF diet were able to lose approximately 30% body weight within 1 week despite maintaining a high calorie intake (Fig. 5e). This weight loss was associated with major reductions in fat mass (Extended Fig. 6d). With weight loss, cysteine deficient mice had improved metabolic homeostasis, (Fig. 5f and Extended data Fig. 6e,f), increased EE (Fig. 5g,h). Notably, immuno-histological analysis of the white adipose depots demonstrated that cysteine deficiency induced browning even while on HFD with increased expression of UCP1 in SFAT and VFAT (Fig. 5i). Furthermore, cysteine-deficiency in obese mice reduced RER suggesting higher fat-utilization (Fig. 5j). Additionally, consistent with improvement of metabolic function in obesity, the gene expression of inflammasome components *Il1b, Il18, Nlrp3, Casp1* and pro-inflammatory cytokines *Il6* and *Tnf* were reduced in F4/80^+^CD11b^+^ adipose tissue macrophages in visceral adipose tissue (Fig. 5k) These results demonstrate that induction of cysteine deficiency can cause weight-loss in mouse model of diet-induced obesity, opening new avenues for future drug development for excess weight-loss

## Discussion

Adipose tissue regulates metabolism by orchestrating inter-organ communication required for healthy longevity^53^. Analyses of adipose tissue of humans that underwent moderate CR in free-living conditions have highlighted genes and pathways that link energy metabolism and inflammation to influence healthspan^5, 7^. In rodents, restriction of calories up to 40% reduces core-body temperature (CBT) and induces browning of the adipose tissue of mice reared in sub-thermoneutral temperature^1^. The CR in humans upregulated the fatty acid oxidation and futile lipid cycling induced-thermogenic pathways but UCP1 was undetectable in adipose tissue of CALERIE-II participants^5^. Similarly, weight loss in obese humans is not associated with classical UCP1 adipose tissue browning^54^. This suggests that alternate UCP1-independent mechanisms maybeat play in human and rodent adipose tissue browning and thermogenesis in response to CR, may be due to extreme CR (>40%) or another phenomenon, including reduction of specific amino acids or macronutrients. In this regard, reduction of core-body temperature^55^ and increased FGF21 is a common link between CR and MR-induced adipose browning and increased longevity^1,2,42^. Our studies demonstrated that reduction of cysteine and subsequent rewiring of downstream cysteine metabolism is linked to adipose browning and weight loss.

Expression and activity of TSP genes CBS and CTH increase when cysteine is low^6^. Indeed, during CR, the TSP is induced to defend against the depletion of cysteine levels. MR regimens that improve lifespan are also restricted or deficient in cysteine^15^, and it is unclear whether methionine or cysteine restriction drives pro-longevity effects. Thus, to understand the metabolic requirement of dietary non-essential amino acid such as cysteine, a genetic mouse model is required that lacks *Cth* in conjunction with restriction of cysteine. Surprisingly, previously reported *Cth* mutant mice originally generated on a 129SvEv mouse strain maintained on cysteine-replete normal chow diet were reported to display hypertension and motor-dysfunction characteristic of neurodegenerative changes in corpus striatum^56,57^. Through conditional deletion of *Cth* (on pure C57/B6 background) in adipose tissue and liver, and rescue of weight-loss by cysteine repletion, our data establishes that systemic cysteine depletion drives adipose tissue thermogenesis without causing behavioral defects or pathological lesions.

While it is still unclear why cysteine deficiency triggers the activation of adipose browning, the mechanism of thermogenesis depends on sympathetic β3-adrenergic signaling and partially requires FGF21 and can be successfully maintained even in the absence of UCP1 and at thermoneutrality. The cysteine-starvation elevated fatty acid lipolysis-esterification cycle genes, while the genes regulating calcium and creatine cycle were not affected. Future studies of specific ablation of UCP1-independent thermogenic genes in *Cth*^−/−^ mice on cysteine-restriction are required to determine the causal pathway. The model of cysteine loss that produces a strong browning response may thus allow the discovery of an alternate UCP1-independent mechanism of adipose tissue thermogenesis.

In healthy humans undergoing CR, consistent with reduced cysteine, glutathione, a major redox regulator, was reduced in adipose tissue. The *Cth* deficient mice on a cysteine-free diet show a decrease in oxidized GSH with a compensatory increase in *Gclc*, *Gss*, and accumulation of γ-glutamyl-peptides. Despite increased oxidative stress, the adipose tissue histology, RNA sequencing, and lipidomic analysis of BAT did not reveal overt ferroptosis in cysteine-depletion induced weight loss. Future studies may reveal cysteine-dependent alternative protective mechanisms that control redox balance and ferroptosis while sustaining UCP1-independent thermogenesis.

Taken together, this study expands our understanding of pathways activated by pro-longevity dietary interventions that confer metabolic adaptation required to maintain tissue homeostasis. Thus, the manipulation of TSP activity to drive adipose tissue browning also has implications for developing interventions that control adiposity and promote longevity. In humans, restriction of methionine and cysteine increased FGF21 and caused a reduction in body weight with improvement of metabolic parameters^58^. Similar to our findings, the metabolic benefits of methionine+cysteine dietary restriction in humans were greater than methionine-restriction alone^58^. Here, based on human dietary restriction studies, and mouse models of cysteine-deficiency, we demonstrate that cysteine is essential for organismal metabolism as its absence triggers adipose browning with progressive weight loss.

## Acknowledgments

We thank all investigators and staff involved in coordinating and executing CALERIE-II clinical trial and Yale comparative medicine pathology core led by Dr Carmen Booth for support with autopsies and histology. We also thank UTSW, Dallas Metabolic Core facility (supported by National Institute of Diabetes and Digestive and Kidney diseases P30DK127984 -NIH NORC program) for bomb calorimetry analysis. AL is a recipient of Gruber and NSF fellowship. The research in Dixit Lab was supported in part by NIH grants AG031797, AG073969, AG068863, P01AG051459.

## Materials and methods

### Human Samples

The participants in this study were part of the CALERIE Phase 2 (Rochon et al., 2011) study which was a multi-center, parallel-group, randomized controlled trial by recruitment of non-obese healthy individuals. 238 adults participated at 3 different locations: Pennington Biomedical Research Center (Baton Rouge, LA), Washington University (St. Louis, MO) and Tufts University (Boston, MA) (NCT00427193). Duke University, (Durham, NC) served as a coordinating center. Participants were randomly assigned to of 25% caloric restriction or ad libitum caloric intake for two years. CR group participants actually reached 14% of CR^5,8^ (Ravussin et al. 2015). Men were between 20 and 50 years old and women were between 20 and 47 years old. Their body mass index (BMI) was between 22.0 and 27.9 kg/m^2^ at the initial visit. Samples were collected at baseline, 1 year, and 2 years of intervention. Abdominal subcutaneous adipose tissue biopsy was performed on a portion of CR group participants and used for RNA-sequencing and metabolomics in this study. All studies were performed under protocol approved by the Pennington institutional review board with informed consent from participants.

### Mice

All mice were on the C57BL/6J (B6) genetic background. *Cth*^−/−^ mice (C57BL/6NTac-Cth^tm1a(EUCOMM)Hmgu/Ieg^) were purchased from the European Mouse Mutant Cell Repository. Breeding these mice to Flipase transgenic mice from Jackson Laboratories generated *Cth^fl/fl^* mice which were crossed to Adipoq-cre and Albumin-cre, purchased from Jackson Laboratories. *Ucp1*^−/−^ and CHOP^−/−^ mice were purchased from Jackson laboratories and crossed to *Cth*^−/−^ mice. *Fgf21*^−/−^ mice were kindly provided by Dr. Steven Kliewer (UT Southwestern) as described previously^41^ and crossed to *Cth*^−/−^ mice. All mice used in this study were housed in specific pathogen-free facilities in ventilated cage racks that deliver HEPA-filtered air to each cage with free access to sterile water through a Hydropac system at Yale School of Medicine. Mice were fed a standard vivarium chow (Harlan 2018s) unless special diet was provided and housed under 12 h light/dark cycles. All experiments and animal use were approved by the Institutional Animal Care and Use Committee (IACUC) at Yale University.

### Diet studies

For cysteine deficiency studies, mice were fed either a control diet, CysF diet, HFD-CTRL diet, or HFD-CysF diet purchased from Dyets, for 6 days unless specified otherwise. For pair feeding studies, mice were provided with either ad libitum or 2.22-2.27g of diet daily.

### Western blot analysis

Cell lysates were prepared using RIPA buffer and optionally frozen and stored at −80°C. Samples were left on ice, vortexing every ten min for 30 min. For tissue samples, snap frozen tissues were ground by mortar and pestle in liquid nitrogen and resuspended in RIPA buffer with protease and phosphatase inhibitors. Samples were centrifuged at 14,000g for 15min and the supernatant was collected protein concentration was determined using the DC Protein Assay (Bio-Rad) and transferred to a nitrocellulose membrane. The following antibodies (and source) were used to measure protein expression: β-Actin (Cell Signaling), pHSL p660 (Cell Signaling), ATGL (Cell Signaling), UCP1 (Abcam), CSE (Novus), Tubulin (Sigma), HSL (Cell Signaling), COMT (Biorad), MAOA (Abcam), TH (Cell Signaling), IRE1a (Cell Signaling), Calnexin (Cell Signaling), BiP (Cell Signaling), CHOP (Cell Signaling), HSP90 (Cell Signaling); followed by incubation with appropriate HRP-conjugated secondary antibodies (Thermo Fisher Scientific).

### Gene expression analysis

Cells or ground tissue (described above) were collected in STAT-60 (Tel-test). RNA from cells were extracted using Qiagen RNeasy micro kits following manufacturer’s instructions. For tissue samples, RNA was extracted using Zymo mini kits following manufacturer’s instructions. During RNA extraction, DNA was digested using RNase free DNase set (Qiagen). Synthesis of cDNA was performed using iScript cDNA synthesis kit (Bio-Rad) and real time quantitative PCR (Q-PCR) was conducted using Power SYBR Green detection reagent (Thermo Fischer Scientific) on a Light Cycler 480 II (Roche).

### Glucose tolerance test

*Cth*^−/−^ HFD-CTRL and HFD-CysF mice were fasted 14hr prior to glucose tolerance test. Glucose was given by i.p. injection based on body weight (0.4g/kg). *Cth*^−/−^ CTRL and CysF mice were fasted for 4hr. Glucose was given by i.p based on lean mass determined by Echo-MRI (2g/kg of lean mass). Blood glucose levels were measured by handheld glucometer (Breeze, Bayer Health Care).

### Flow Cytometry

Adipose tissue was digested at 37°C in HBSS (Life Technologies) + 0.1% collagenase I or II (Worthington Biochemicals). The stromal vascular fraction was collected by centrifugation, washed and filtered using 100um and 70um strainers. Cells were stained with LIVE/DEAD™ Fixable Aqua Dead Cell Stain Kit (Thermo Fisher Scientific) and then for surface markers including CD45, CD3, B220, CD11b, F4/80, Ly6G, Siglec F, CD163, CD24, F3, CD31, Pdgfra, Dpp4, and CD9 and all antibodies were purchased from eBioscience or Biolegend. Cells were fixed in 2% PFA. Samples were acquired on a custom LSR II and data was analyzed in FlowJo.

### Single-cell RNA sequencing

For stromal vascular fraction, female *Cth*^+/+^ and *Cth*^−/−^ mice were fed CTRL of CysF diet for 4 days. SFAT was collected, with lymph nodes removed, pooled, and digested. Isolated cells were subjected to droplet-based 3’ end massively parallel single-cell RNA sequencing using Chromium Single Cell 3’ Reagent Kits as per manufacturer’s instructions (10x Genomics). The libraries were sequenced using a HiSeq3000 instrument (Illumina). Sample demultiplexing, barcode processing, and single-cell 3’ counting was performed using the Cell Ranger Single-Cell Software Suite (10x Genomics). Cellranger count was used to align samples to the reference genome (mm10), quantify reads, and filter reads with a quality score below 30. The Seurat package in R was used for subsequent analysis^31^. Cells with mitochondrial content greater than 0.05% were removed and data was normalized using a scaling factor of 10,000, and nUMI was regressed with a negative binomial model. Principal component analysis was performed using the top 3000 most variable genes and t-SNE analysis was performed with the top 20 PCAs. Clustering was performed using a resolution of 0.4. The highly variable genes were selected using the FindVariableFeatures function with mean greater than 0.0125 or less then 3 and dispersion greater than 0.5. These genes are used in performing the linear dimensionality reduction. Principal component analysis was performed prior to clustering and the first 20 PC’s were used based on the ElbowPlot. Clustering was performed using the FindClusters function which works on K-nearest neighbor (KNN) graph model with the granularity ranging from 0.1-0.9 and selected 0.4 for the downstream clustering. For identifying the biomarkers for each cluster, we have performed differential expression between each cluster to all other clusters identifying positive markers for that cluster. To understand the trajectory of the adipocyte progenitors, we used Monocle2 to analyze scRNA-seq data of Clusters 0, 1, and 2 (Trapnell 2014).

### Whole tissue RNA sequencing and transcriptome analysis

Snap frozen tissues were ground by mortar and pestle in liquid nitrogen and resuspended in STAT-60. RNA was extracted using Zymo mini kits. RNA was sequenced on a HiSeq2500. The quality of raw reads was assessed with FastQC [FastQC]. Raw reads were mapped to the GENCODE vM9 mouse reference genome [GENCODE] using STAR aligner [STAR] with the following options: --outFilterMultimapNmax 15 --outFilterMismatchNmax 6 --outSAMstrandField All --outSAMtype BAM SortedByCoordinate --quantMode TranscriptomeSAM. The quality control of mapped reads was performed using in-house scripts that employ Picard tools [Picard]^5^. The list of rRNA genomic intervals that we used for this quality control was prepared on the basis of UCSC mm10 rRNA annotation file [UCSC] and GENCODE primary assembly annotation for vM9 [GENCODE]. rRNA intervals from these two annotations were combined and merged to obtain the final list of rRNA intervals. These intervals were used for the calculation of the percentage of reads mapped to rRNA genomic loci. Strand specificity of the RNA-Seq experiment was determined using an in-house script, on the basis of Picard [Picard] mapping statistics. Expression quantification was performed using RSEM [RSEM]. For the assessment of expression of mitochondrial genes, we used all genes annotated on the mitochondrial chromosome in the GENCODE vM9 mouse reference genome [GENCODE]. PCA was performed in R. For the PCA, donor effect was removed using the ComBat function from the sva R-package [sva]. Gene differential expression was calculated using DESeq2 [DESeq2].]. Pathway analysis was done using fgsea (fast GSEA) R-package [fgsea] with the minimum of 15 and maximum of 500 genes in a pathway and with 1 million of permutations. For the pathway analysis, we used the Canonical Pathways from the MSigDB C2 pathway set [MSigDB1, MSigDB2], v6.1. The elimination of redundant significantly regulated pathways (adjusted p-value < 0.05) was done using an in-house Python script in the following way. We considered all ordered pairs of pathways, where the first pathway had normalized enrichment score equal to or greater than the second pathway. For each ordered pair of pathways, we analyzed the leading gene sets of these pathways. The leading gene sets were obtained using fgsea [fgsea]. If at least one of the leading gene sets in a pair of pathways had more than 60% of genes in common with the other leading gene set, then we eliminated the second pathway in the pair.

### Sample preparation for metabolome analysis

Frozen tissues or serum samples, together with internal standard compounds (mentioned below), was subjected to sonication in 500μL of ice-cold methanol. To this, an equal volume of ultrapure water (LC/MS grade, Wako, Japan) and 0.4 volume of chloroform were added. The resulting suspension was centrifuged at 15,000×g for 15 minutes at 4 °C. The aqueous phase was then filtered using an ultrafiltration tube (Ultrafree MC-PLHCC, Human Metabolome Technologies, Japan), and the filtrate was concentrated by nitrogen spraying (aluminum block bath with nitrogen gas spraying system, DTU-1BN/EN1-36, TAITEC, Japan). The concentrated filtrate was dissolved in 50μL of ultrapure water and utilized for IC-MS and LC-MS/MS analysis. Methionine sulfone and 2-morpholinoethanesulfonic acid were employed as internal standards for cationic and anionic metabolites, respectively. The recovery rate (%) of the standards in each sample measurement was calculated to correct for the loss of endogenous metabolites during sample preparation.

### IC-MS metabolome analysis

Anionic metabolites were detected using an orbitrap-type MS (Q-Exactive focus; Thermo Fisher Scientific, USA) connected to a high-performance ion-chromatography (IC) system (ICS-5000+, Thermo Fisher Scientific, USA) that allows for highly selective and sensitive metabolite quantification through IC separation and Fourier transfer MS principle. The IC system included a modified Thermo Scientific Dionex AERS 500 anion electrolytic suppressor, which converted the potassium hydroxide gradient into pure water before the sample entered the mass spectrometer. Separation was carried out using a Thermo Scientific Dionex IonPac AS11-HC column with a particle size of 4μm. The IC flow rate was 0.25 mL/min, supplemented post-column with a makeup flow of 0.18 mL/min MeOH. The potassium hydroxide gradient conditions for IC separation were as follows: from 1 mM to 100 mM (0–40 min), to 100 mM (40–50 min), and to 1 mM (50.1–60 min), with a column temperature of 30 °C. The Q Exactive focus mass spectrometer was operated in the ESI-negative mode for all detections. A full mass scan (m/z 70–900) was performed at a resolution of 70,000. The automatic gain control target was set at 3×10^6^ ions, and the maximum ion injection time was 100ms. The source ionization parameters were optimized with a spray voltage of 3 kV, and other parameters were as follows: transfer temperature, 320 °C; S-Lens level = 50, heater temperature, 300 °C; sheath gas = 36, and Aux gas, 10.

### LC-MS/MS metabolome analysis

Cationic metabolites were measured using liquid chromatography-tandem mass spectrometry (LC-MS/MS). The LCMS-8060 triple-quadrupole mass spectrometer (Shimadzu corporation, Japan) with an electrospray ionization (ESI) ion source was employed to perform multiple reaction monitoring (MRM) in positive and negative ESI modes. The samples were separated on a Discovery HS F5-3 column (2.1 mm I.D. x 150 mm L, 3μm particle, Sigma-Aldrich) using a step gradient of mobile phase A (0.1% formate) and mobile phase B (0.1% acetonitrile) with varying ratios: 100:0 (0-5 min), 75:25 (5-11 min), 65:35 (11-15 min), 5:95 (15-20 min), and 100:0 (20-25 min). The flow rate was set at 0.25 mL/min, and the column temperature was maintained at 40°C.

### Monoamine measurements by HPLC with electro chemical detector (ECD)

For low concentration monoamine measurements, extracted tissue metabolites by abovementioned protocol were injected with an autosampler (M-510, Eicom) into a HPLC unit (Eicom) coupled to an ECD (ECD-300, Eicom). The samples were resolved on the Eicompak SC-5ODS column (φ3.0 x 150 mm, Eicom), using an isocratic mobile phase (5 mg/L EDTA-2Na, 220 mg/L sodium 1-octanesulfonate in acetate/citrate buffer (0.1 M, pH 3.5)/MeOH (83:17, v/v)), at a flow rate of 0.5 mL/min and a column temperature of 25°C. At the ECD, analytes were subjected to oxidation reactions within the ECD unit with WE-3G graphite electrode (applied potential is +750 mV against an Ag/AgCl reference electrode). Resulting chromatograms were analyzed using the software EPC-300 (Eicom).

### Lipidome analysis

To extract total lipids, frozen tissues were mixed with 500 μL of 1-butanol/methanol (1:1, v/v) containing 5 mM ammonium formate. The mixture was vortexed for 10 seconds, sonicated for 15 minutes in a sonic water bath, and then centrifuged at 16,000 × g for 10 minutes at 20°C. The supernatant was transferred to a 0.2-mL glass insert with a Teflon insert cap for LC ESI-MS analysis.

For lipidomic analysis, a Q-Exactive focus orbitrap mass spectrometer (Thermo Fisher Scientific, San Jose, CA) was connected to an HPLC system (Ultimate3000, Thermo Fisher Scientific). The samples were separated on a Thermo Scientific Accucore C18 column (2.1 × 150 mm, 2.6 μm) using a step gradient of mobile phase A (10 mM ammonium formate in 50% acetonitrile and 0.1% formic acid) and mobile phase B (2 mM ammonium formate in acetonitrile/isopropyl alcohol/water, ratios of 10:88:2, v/v/v, with 0.02% formic acid). The gradient ratios used were 65:35 (0 min), 40:60 (0-4 min), 15:85 (4-12 min), 0:100 (12-21 min), 0:100 (21-24 min), 65:35 (24-24.1 min), and 100:0 (24.1-28 min) at a flow rate of 0.4 mL/min and a column temperature of 35°C.

The Q-Exactive focus mass spectrometer operated in both positive and negative ESI modes. It performed a full mass scan (m/z 250-1100), followed by three rapid data-dependent MS/MS scans, at resolutions of 70,000 and 17,500, respectively. The automatic gain control target was set at 1 × 10^6^ ions, and the maximum ion injection time was 100 ms. The source ionization parameters included a spray voltage of 3 kV, transfer tube temperature of 285°C, S-Lens level of 45, heater temperature of 370°C, sheath gas at 60, and auxiliary gas at 20. The acquired data were analyzed using LipidSearch software (Mitsui Knowledge Industry, Tokyo, Japan) for major phospholipids (PLs). The search parameters for LipidSearch software were as follows: precursor mass tolerance = 3 ppm, product mass tolerance = 7 ppm, and m-score threshold = 3.

### Visualizing noradrenaline distribution using MALDI-imaging mass spectrometry

The tissue block was frozen and secured onto a disc using a cryoembedding medium (Super Cryoembedding Medium, SECTION-LAB, Hiroshima, Japan), then equilibrated at −16°C in cryostats (Leica Biosystems, Nussloch, Germany). Tissue sections, 8 µm thick, were cut and mounted onto conductive indium-tin-oxide (ITO)-coated glass slides (Matsunami Glass Industries, Osaka, Japan)., A solution of tetrafluoroborate salts of 2,4-diphenyl-pyrylium (DPP) (1.3 mg/mL in methanol) for on-tissue derivatization of monoamines, and DHB-matrix (50 mg/mL in 80% ethanol) were manually sprayed onto the tissue using an airbrush (Procon Boy FWA platinum; Mr. Hobby, Tokyo). The manual spray was performed at room temperature, applying 40 μL/mm2 with a distance of approximately 50 mm. The samples were analyzed using a linear ion trap mass spectrometer (LTQ XL, Thermo Fisher Scientific). The raster scan pitch was set at 50 µm. Signals of noradrenaline-DPP (m/z 384 > 232) were monitored with a precursor ion isolation width of m/z 1.0 and a normalized collision energy of 45%. Ion images were reconstructed using ImageQuest 1.1.0 software (Thermo Fisher Scientific).

### Core-body temperature measurement

Animals were anesthetized with isoflurane, first at a rate of 2-3% and maintained at 0.5-2% in oxygen during surgery. Mice were kept on a heating pad throughout surgery. Mice were injected with buprenorphine and bupivacaine as pre-emptive analgesia. A small ventral incision of 1cm was made after clipping hair and disinfection with betadine and 70% ethanol. DST nano-T temperature loggers (Star Oddi) were placed in the peritoneal cavity, and abdominal muscle and skin were sutured closed. Post-surgery, mice were singly housed and provided with Meloixcam for 48 hours. After 7 days, sutures were removed. 10 days after surgery, mice were started on CTRL or CysF diet, and loggers were removed for data collection after euthanization. Loggers were programmed to take temperature readings every 30 minutes.

### Metabolic cages

The energy expenditure (EE), respiratory exchange ratio (RER), activity, food intake of mice were monitored using the TSE PhenoMaster System (V3.0.3) Indirect Calorimetry System. Each mouse was housed in individual chambers for 3 days for acclimation and switched to experimental diet for 6 days. Each parameter was measured every 30 min. EE and RER were calculated based on the oxygen consumption (O_2_) and carbon dioxide production (CO_2_). Mouse activity was detected by infrared sensors, and food intake and water consumption were measured via weight sensors on food and water dispensers located in the cage.

### EchoMRI

The parameters of body composition were measured in vivo by magnetic resonance imaging (EchoMRI; Echo Medical Systems). The amount of fat mass, lean mass and free water were measured by the analysis. For the analysis, each mouse was placed in an acrylic tube with breathing holes and the tube was inserted in the MRI machine. The analysis per mouse takes approximately 90 sec and automatically calculated numerical results were analyzed.

### Climate chambers

Mice were acclimated in climate chambers (model 7000-10, Caron) at either 30°C or 20°C, with humidity maintained at 50% under 12 h light/dark cycles. After one week acclimation, mice were switched to either CTRL or CysF diet for 6 days, while maintained in the climate chambers. Mice were handled daily to measure body weight.

### Feces bomb calorimetry

Feces were collected daily over the course of CTRL or CysF feeding. Samples were dried for 72 hours. Fecal bomb calorimetry was performed at UT Southwestern Medical Center Metabolic Phenotyping Core (Dallas, TX, USA) using a Parr 6200 Isoperibol Calorimeter equipped with a 6510 Parr Water Handling.

### Serum measurements

After blood collection by cardiac puncture, samples were allowed to clot for 2 hours. Serum was collected after centrifugation. FGF21 and GDF15 levels in the serum were measured by ELISA (R&D). Cysteine levels were determined by competitive EIA (LS-Bio). Glycerol levels were determined by colorimetric assay (Sigma Aldrich).

### β-3 adrenergic receptor inhibition

Mice were administered twice daily L748337 (Santa Cruz Biotechnology) (5mg/kg) by i.p injection. Mice were weighed daily and assessed for their health.

### Histology

Tissues were collected in 10% formalin, embedded in paraffin and sectioned into 5um thick sections. Tissues were stained with hematoxylin and eosin (H&E) or stained for UCP1 (Abcam) and Goat anti-rabbit HRP (DAKO) and developed for color using Abcam DAB substrate kit.

### Animal preparation for BIRDS Temperature Analysis

The animals were anesthetized with 3% isoflurane in an induction chamber and then kept at 2-3% during surgery. The animal was laid back on a microwaveable heating pad. Prior to incision, a single dose of bupivacaine was given for analgesia. A 1-2 cm midline incision was made on the neck to expose the jugular vein. Another small incision (<1 cm) was made at the back of the neck. A sterile polyurethane or silicone catheter with a metal guide was inserted from the back of the neck, where the vascular port was fixed to the jugular vein. Prior to implantation the port and the catheter were flushed with heparinized saline (25 IU/ml). The jugular vein was catheterized toward the heart. The skin was closed with surgical sutures after application of triple antibiotic ointment and the vascular port was fixed. The duration of the surgical procedure was 15-20 min.

### MR data acquisition

TmDOTMA^−^ was purchased from Macrocyclics (Plano, TX, USA). Temperature mapping with BIRDS was performed on a 9.4T Bruker scanner (Billerica, MA). The respiration rate was monitored during the entire duration of the experiment. A 200mM TmDOTMA^−^ solution was infused at a rate of 60 to 80 µl/h for 1 to 2 hours. The infusion rate was adjusted according to animal physiology. The T_2_ weighted magnetic resonance (MR) images were acquired with an FOV of 23×23mm^2^, 128×128 matrix, 23 slices of 0.5mm thickness, TR=3s and TE=9ms. The extremely short T_1_ and T_2_ relaxation times (<5ms) of the TmDOTMA^−^ methyl group allowed ultrafast temperature mapping with BIRDS using 3D chemical shift imaging (CSI) acquisition with a short TR (10ms) and wide bandwidths (±150ppm). Temperature mapping with BIRDS was started immediately after detection of global MR signal of TmDOTMA^−^ methyl group, at about 1 hour after the start of the infusion. The CSI was acquired using a FOV of 23×15×23mm^3^, 809 spherical encoding steps, 21min acquisition, and reconstructed to 23×15×23, with a voxel resolution of 1×1×1mm^3^. Selective excitation of the TmDOTMA^−^ methyl group was achieved using a single band 200µs Shinnar-Le Roux (SLR) RF pulse. The MR spectrum in each voxel was line broadened (200 Hz) and phased (zero-order) in Matlab (MathWorks Inc., MA, USA), and the corresponding temperature T_c_ was calculated from the chemical shift δ_CH3_ of the TmDOTMA^−^ methyl group according to

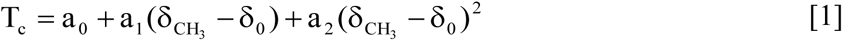

where δ_0_ = −103.0 ppm and the coefficients a_0_ = 34.45 ± 0.01, a_1_ = 1.460 ± 0.003 and a_2_ = 0.0152 ± 0.0009 were calculated from the linear least-squares fit of temperature as a function of chemical shift δ_CH3_ (reference below). Statistical analysis was done using Student’s t-test with two tails, with p<0.05 used as a cutoff for significance.

### In vivo spin trapping and Electron Paramagnetic Resonance (EPR) spectroscopy

POBN (α-(4-Pyridyl-1-oxide)-N-t-butylnitrone, Enzo) was used for spin trapping; POBN was dissolved in saline and administered i.p. at 500 mg/kg body weight. Tissue samples (VFAT, SFAT and BAT) were collected 45 minutes post-injection, immediately frozen in liquid nitrogen, and stored at −80°C until EPR measurements. Lipid extraction was performed using chloroform/methanol (2/1) (Folch-extraction) as described previously^59^. All EPR spectra were recorded in a quartz flat cell using an X-band EMX plus EPR Spectroscope (parameters: 3,480 ± 80 G scan width, 105 receiver gain, and 20 mW microwave power; time constant: 1,310 ms; conversion time: 655 ms).

### Aconitase activity

Aconitase activity was measured with Aconitase Assay kit (Cayman). Freshly collected SFAT and VFAT samples were measured at 500 μg total protein/mL, and BAT samples were measured at 100 μg total protein/mL. All results were normalized to 500 μg/mL total protein concentration. Standard protocols provided with the kits was followed.

### *In vitro* adipocyte differentiation

Stromal vascular fraction from visceral depots of *Cth*^−/−^ was isolated as previously described. Cells were plated in growth medium (DMEM supplemented with 10% FBS and 1% Penicillin-Streptomycin) and expanded for 3-5 days. Adipocyte differentiation was induced with growth medium supplemented with insulin (5μg/ml), rosiglitazone (1μM), iso-butyl-methylxanthine (0.5mM) and dexamethasone (1μM) for 48hrs. Cells were maintained on differentiation medium containing insulin (5μg/ml) and rosiglitazone (1μM) for 96hr. Fully differentiated cells were then treated with various concentrations of Cystine (0-200μM) for 48hr, in cystine and methionine-free DMEM (Gibco) supplemented with 10% dialyzed FBS, 1% Penicillin-Streptomycin and 200μM methionine.

### Quantification and statistical analysis

Statistical differences between groups were calculated by unpaired t-tests. For comparing groups over time, mice were individually tracked and groups were compared using 2-way ANOVA with Sidak’s correction for multiple comparisons. For all experiments a p-value of p≤0.05 was considered significant.

---

**Extended Data Figure 1:**
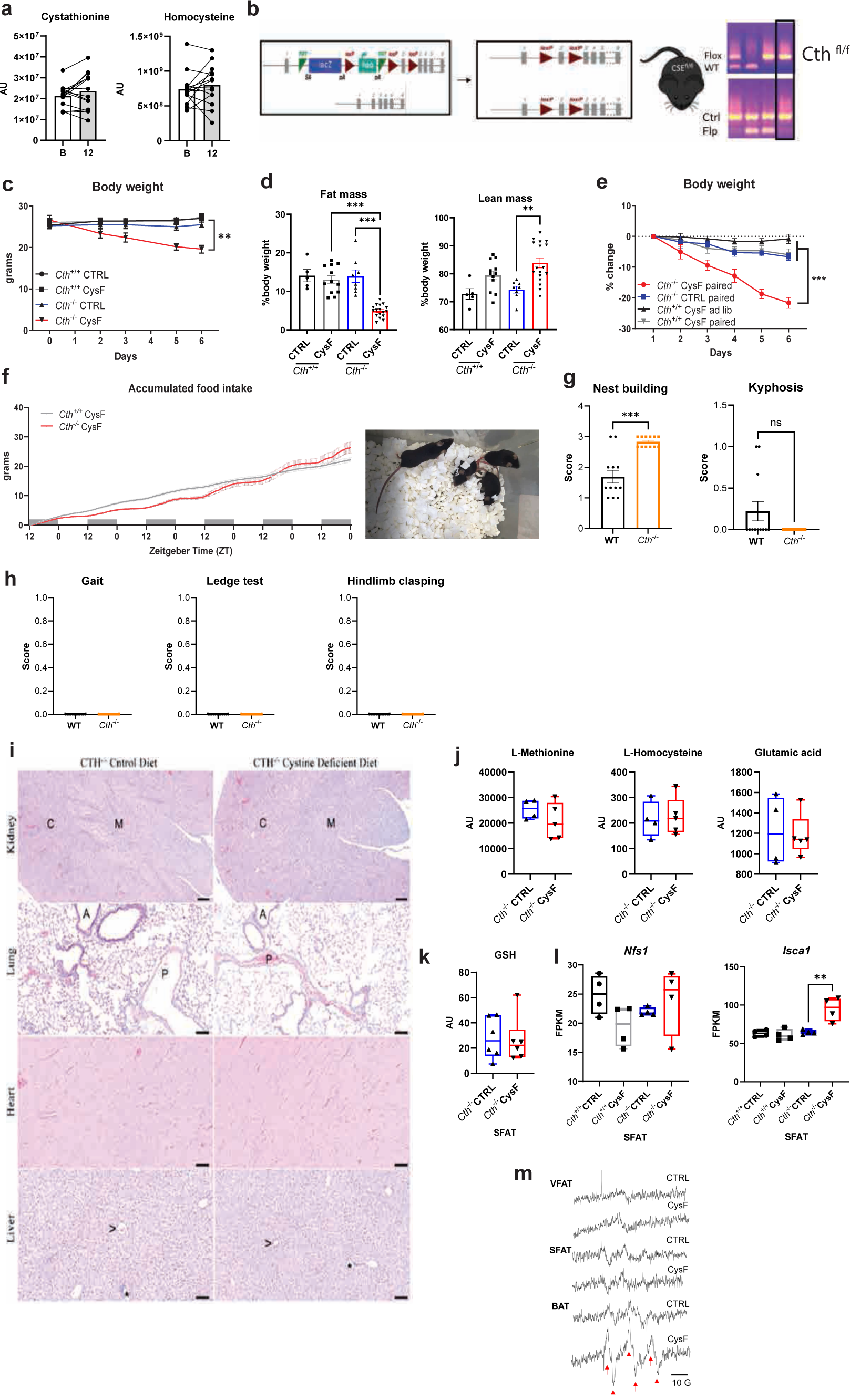
Cysteine depletion induces weight-loss in mice without overt pathology. a) Cystathionine and homocysteine measurements by MS/MS in human SFAT at baseline (B) and after 12 months of caloric restriction (n=14). AU: arbitrary units. b) Schematic of *Cth^−/−^* and *Cth*^fl/fl^ mice generation (KOMP construct) used to cross to either Alb:cre or Adipoq:cre. c) Body weight of *Cth*^+/+^ and *Cth*^−/−^ mice fed with CTRL or CysF diet for 6 days (n=5 *Cth*^+/+^ CTRL, n=6 *Cth*^+/+^ CysF, n=4 *Cth*^−/−^ CTRL, n=5 *Cth*^−/−^ CysF). d) Fat mass and lean mass measured by EchoMRI of male *Cth*^+/+^ and *Cth*^−/−^ after 6 days of CTRL or CysF diet (n=5 *Cth*^+/+^ CTRL, n=12 *Cth*^+/+^ CysF, n=8 *Cth*^−/−^ CTRL, n=17 *Cth*^−/−^ CysF). e) *Cth*^+/+^ and *Cth*^−/−^ mice were fed ad libitum (ad lib) or pair fed CTRL or CysF diet (n=4 *Cth*^+/+^CysF ad lib, n=5 *Cth*^+/+^CysF pair fed, n=7 *Cth*^−/−^ CTRL pair fed, n=5 *Cth*^−/−^ CysF pair fed). Percentage body weight change over 6 days of diet. f) Accumulated food intake of *Cth*^+/+^ and *Cth*^−/−^ mice over 6 days of CysF feeding measured in metabolic cages (n=10 *Cth*^+/+^ and n=12 *Cth*^−/−^). Cage image and video show that *Cth*^−/−^ mice on CysF diet at day 5 have normal activity. g) Qualitative assessment of nest building (score from 0 to 4) and presence (score=1) or absence (score=0) of kyphosis in WT and *Cth^−/−^* mice (n=12/group). h) Gait assessment, ledge test and hindlimb clasping test were performed to measure motor coordination in WT and *Cth*^−/−^ mice. Mice were scored from 0 (normal behavior) to 1 (abnormal behavior) (n=12/group). i) Representative H&E-stained sections of kidney, lung, heart, and liver from female CTH^−/−^ mice fed control diet or Cystine-deficient diet for 6 days, lack significant pathologic changes and do not differ in microscopic changes by diet in the tissues examined. C = renal cortex, M = renal medulla A = airway, P = pulmonary artery, > = central vein, and * = portal triad. Kidney scale bars=200 μm, lung, heart, liver scale bars= 100μm. j) Serum L-methionine, L-homocysteine, glutamic acid and k) SFAT GSH quantified by mass spectrometry in *Cth*^−/−^ mice fed with CTRL or CysF diet for 6 days (n=4-5/group). AU: arbitrary units. l) RNA-seq-based *Nfs1* and *Isca1* gene expression in SFAT of *Cth*^+/+^ and *Cth*^−/−^ mice after 6 days of CTRL or CysF feeding (n=4/group). m) Representative EPR spectra of POBN-lipid radical adducts measured in Folch extracts of VFAT, SFAT and BAT tissues. The six-line spectrum (red arrows) is consistent with carbon-centered lipid-derived radicals, indicative of lipid peroxidation (identified through hyperfine coupling constants a^N^ = 15.75 ± 0.06 G and *a_β_*^H^ = 2.77 ± 0.07 G). Data are expressed as mean±SEM. Statistical differences were calculated by 2-way ANOVA with Sidak’s correction for multiple comparisons, or by unpaired t-test (**p<0.01, ***p<0.001).

**Extended Data Figure 2:**
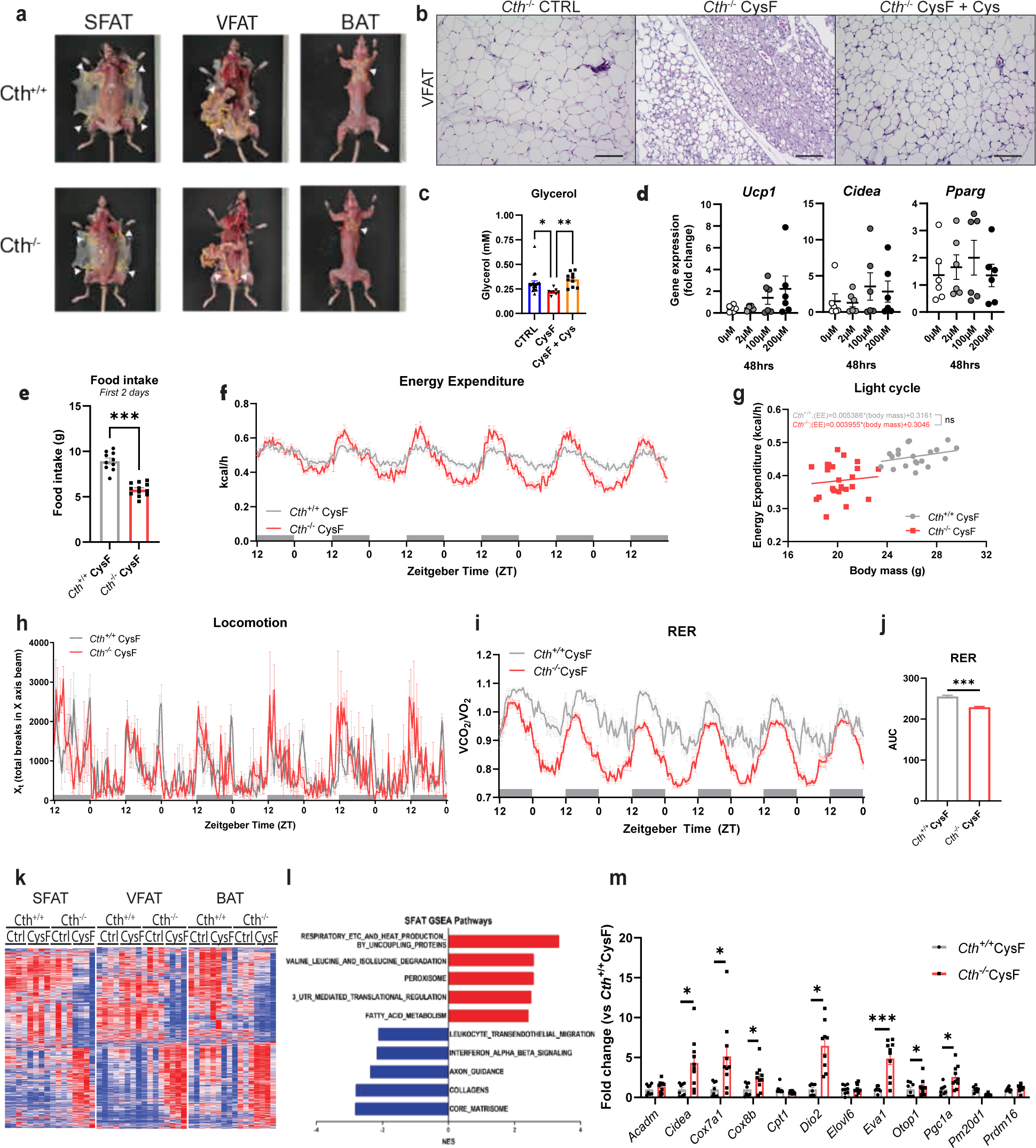
Cysteine starvation induces thermogenic reprogramming of adipose tissue transcriptome. a) Representative subcutaneous (SFAT), visceral (VFAT), and brown adipose depots (BAT) of *Cth*^+/+^ and *Cth*^−/−^ after 6 days of CysF diet. b) Representative H&E-stained sections of VFAT of *Cth*^−/−^ mice fed CTRL or CysF diet for 6 days or after Cys supplementation following CysF weight loss (scale bar=100 μm). c) Serum glycerol levels of *Cth*^−/−^ mice fed with CTRL (n=20) or CysF (n=8) or switched to Cys-containing diet after CysF feeding (n=10). d) *Ucp1*, *Cidea* and *Pparg* gene expression in *Cth*^−/−^ pre-adipocytes differentiated *in vitro* and treated with increasing concentration of Cystine for 48 hours. e) Cumulative food intake during the initial two days of CysF feeding in *Cth*^+/+^ and *Cth*^−/−^ mice (n=10 *Cth*^+/+^ and n=12 *Cth*^−/−^). f-j) *Cth*^+/+^ and *Cth*^−/−^ mice were fed with CysF diet for 6 days and housed in metabolic cages (n=10 *Cth*^+/+^ and n=12 *Cth*^−/−^). f) Energy expenditure during CysF feeding. g) Linear regression analysis of unnormalized average energy expenditure measured by indirect calorimetry against body mass on days 4 and 5 of CysF diet. h) Locomotor activity. i) Respiratory exchange ratio (RER) and j) area under the curve (AUC) quantified for RER. k-l) Whole tissue RNA-seq of SFAT, VFAT, and BAT of *Cth*^+/+^ and *Cth*^−/−^ fed 6 days of CTRL or CysF diet (n=4/group). k) Heat map highlighting changes specifically occurring in cysteine deficiency. l) Select top pathways being up- and down-regulated in *Cth*^−/−^ CysF vs CTRL in SFAT after gene set enrichment analysis. i) Gene expression of selected thermogenesis markers confirmed by qPCR in SFAT, in *Cth*^+/+^ and *Cth*^−/−^ mice fed with CysF diet (n=8 *Cth^+/+^* and n=10 *Cth^−/−^*. Data are expressed as mean±SEM. Statistical differences were calculated by one-way ANOVA, or by 2-way ANOVA with Sidak’s correction for multiple comparisons, or by unpaired t-test, (*p<0.05, **p<0.01, ***p<0.001).

**Extended Data Figure 3:**
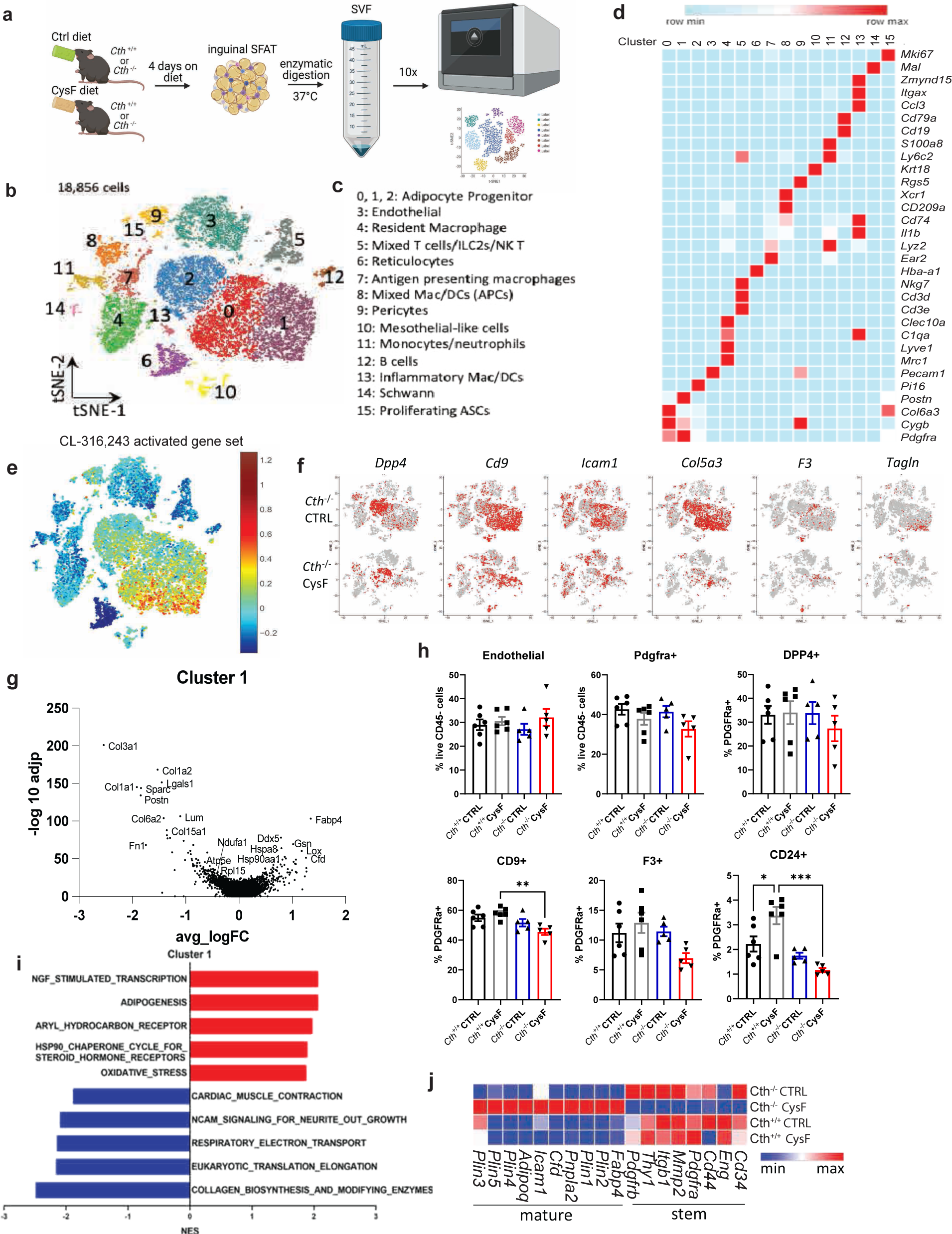
Impact of cysteine depletion on transcriptional regulation of adipose tissue at single cell resolution. a) Experimental design schematic of cell processing of subcutaneous adipose depot (SFAT) stromal vascular fraction (SVF) for scRNA-seq. b) t-SNE plot of scRNAseq from SFAT stromal vascular fraction with c) cluster identities. APCs: antigen presenting cells. ASCs: adipose-derived stromal cells. d) Heat map of normalized gene expression of selected markers to identify major cell lineages. e) Enrichment of CL-316,243 activated gene signature overlaid on all populations in all samples. f) t-SNE plots displaying *Dpp4*, *Cd9*, *Icam1*, *Col5a3*, *F3*, and *Tagln* expression in red across all populations in *Cth*^−/−^ CTRL and *Cth*^−/−^ CysF samples. g) Volcano plot of differentially expressed genes comparing *Cth*^−/−^ CysF and *Cth*^+/+^ CysF in cluster 1. h) Orthogonal validation of adipocyte progenitor changes using FACS analysis of SFAT SVF in *Cth*^+/+^ and *Cth*^−/−^ mice on CTRL and CysF diet for 4 days (n=5-6/group). i) Select top pathways from gene set enrichment comparing *Cth*^−/−^ CysF vs. *Cth*^+/+^ CysF in cluster 1. j) Heatmap of gene expression of select stem and mature adipocyte genes in clusters 0, 1 and 2 showing the impact of cysteine depletion in mice. Data are expressed as mean±SEM. Statistical differences were calculated by 2-way ANOVA with Sidak’s correction for multiple comparisons, and by unpaired t-test (*p<0.05, **p<0.01, ***p<0.001).

**Extended Data Figure 4:**
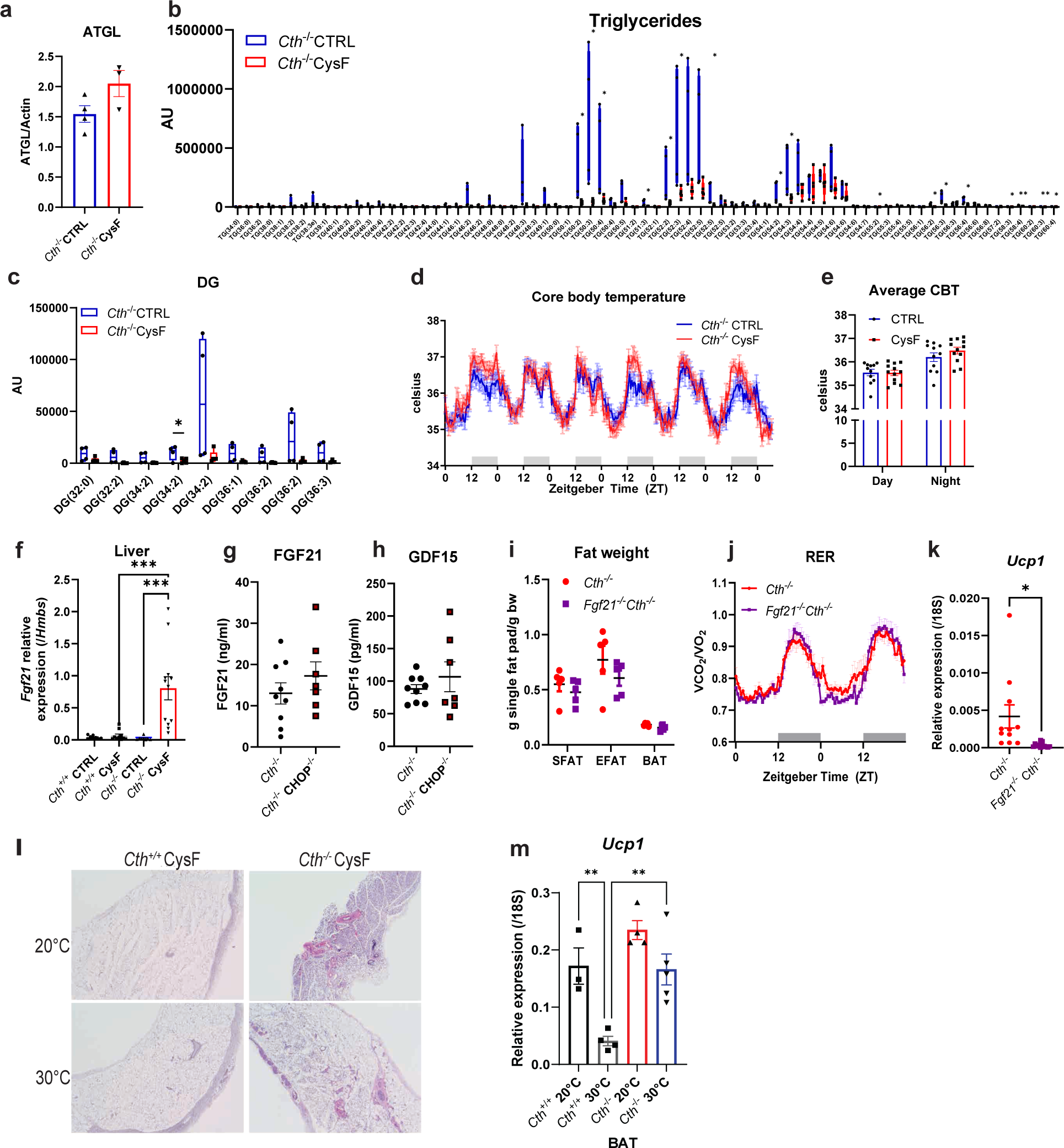
Cysteine-depletion mobilizes lipids for thermogenic response independently of thermoneutrality. a) Quantification of ATGL immunoblot shown in Fig. 3a, Actin was used as a loading control. b-c) Tissue lipidomics of brown adipose depot (BAT) from *Cth*^−/−^ mice fed CTRL (n=4) or CysF diet (n=5) for 6 days with b) triglycerides (TG) and c) diacylglycerol species highlighted. AU: arbitrary units. d) Core body temperature (CBT) measured in the peritoneal cavity by implantation of Star-Oddi logger of *Cth*^−/−^ mice fed with CTRL or CysF diet over 6 days and e) average day and night CBT of *Cth*^−/−^ mice fed with CTRL or CysF diet. Recordings were taken every 30 minutes (n=11 *Cth*^−/−^ CTRL, n=12 *Cth*^−/−^ CysF, 3 independent experiments pooled). f) *Fgf21* gene expression in the liver of *Cth*^+/+^ and *Cth*^−/−^ mice fed CTRL or CysF diet for 6 days (n=8 *Cth*^+/+^ CTRL, n=10 *Cth*^+/+^ CysF, n=8 *Cth*^−/−^ CTRL, n=12 *Cth*^−/−^ CysF). g-h) Serum levels of g) FGF21 and h) GDF15 in *Cth*^−/−^ and *Cth*^−/−^CHOP^−/−^ mice after 5 days of CysF feeding, measured by ELISA (n=9 *Cth*^−/−^ and n=7 *Cth*^−/−^CHOP^−/−^). i) SFAT, VFAT and BAT weight normalized to body weight of *Cth*^−/−^ and *Fgf21*^−/−^*Cth*^−/−^ mice after CysF feeding (n=5/group). j) Respiratory exchange ratio (RER) of *Cth*^−/−^ and *Fgf21*^−/−^*Cth*^−/−^ mice upon CysF feeding, measured at day 3 and 4 in metabolic cages (n=5/group). k) *Ucp1* gene expression in SFAT of *Cth*^−/−^ and *Fgf21*^−/−^*Cth*^−/−^ mice after 6 days of CysF feeding (n=11-12/group).l) Representative H&E histology images of SFAT showing increased browning at day 6 in *Cth*^+/+^ and *Cth*^−/−^ mice fed CysF diet and housed at 20°C or at 30°C. m) *Ucp1* gene expression measured by qPCR in BAT of *Cth*^+/+^ and *Cth*^−/−^ mice fed CysF diet and housed at 20°C or at 30°C for 6 days (n=3-5/group). Data are expressed as mean±SEM. Statistical differences were calculated by 2-way ANOVA with Sidak’s correction for multiple comparisons, and by unpaired t-test (*p<0.05, **p<0.01, ***p<0.001).

**Extended Data Figure 5:**
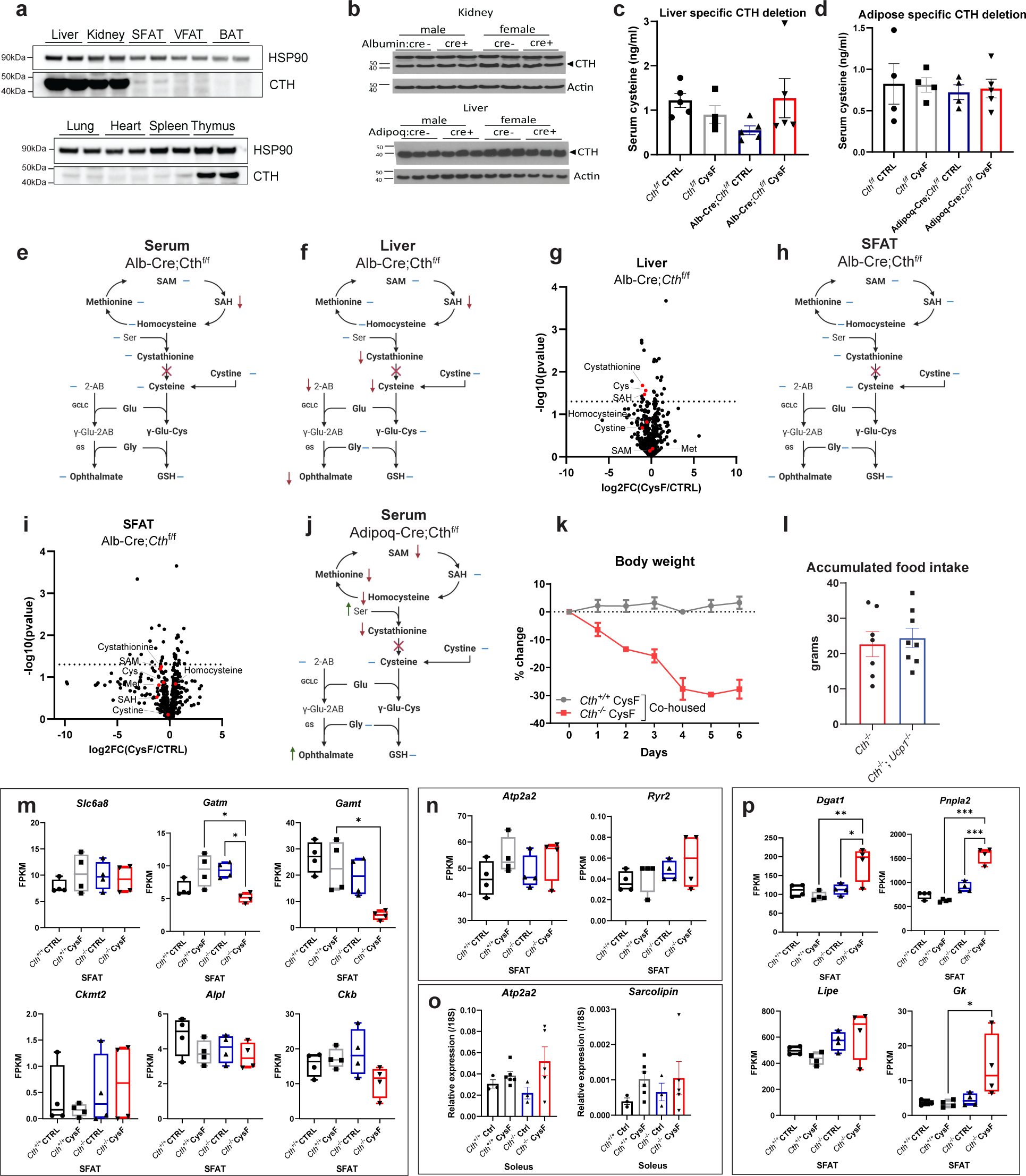
Systemic cysteine depletion induced weight-loss is independent of microbiota and canonical thermogenic pathways. a) Immunoblot analysis of CTH in liver, kidney, subcutaneous (SFAT), visceral (VFAT), brown (BAT) adipose depots, lung, heart, spleen, and thymus. b) Immunoblot analysis of CTH in kidney samples from male and female *Cth*^f/f^;Alb-Cre- and *Cth*^f/f^;Alb-Cre+ mice and in liver samples from male and female *Cth*^f/f^;Adipoq-Cre- and *Cth*^f/f^;Adipoq-Cre+ mice. Actin is used as a loading control. c-d) Cysteine serum levels of c) *Cth*^f/f^ and Alb-Cre;*Cth*^f/f^ mice and d) *Cth*^f/f^ and Adipoq-Cre;*Cth*^f/f^ mice after 5 days of CTRL or CysF diet (n=4-5/group). e-i) Alb-Cre;*Cth*^f/f^ mice were fed CTRL or CysF diet for 6 days. Schematic summary of changes in the metabolites in the e) serum and in the f) liver. g) Volcano plot of metabolites identified by MS/MS in the liver. h) Schematic summary of changes in the metabolites and i) volcano plot of metabolites identified by MS/MS in the SFAT. Transsulfuration pathway related metabolites are highlighted in red. Cys: cysteine. Met: methionine. SAM: S-adenosyl methionine. SAH: S-adenosyl homocysteine. j) Schematic summary of changes in serum metabolites of Adipoq-Cre;*Cth*^f/f^ fed with CTRL or CysF diet for 6 days. Blue lines represent measured, but unchanged metabolites, red and green arrows indicate significantly decreased or increased metabolites, respectively (p<0.05). k) Percentage body weight change of *Cth^+/+^* and *Cth*^−/−^ mice that were co-housed and fed CysF diet for 6 days (n=4/group). l) Accumulated food intake of *Cth*^−/−^ and *Cth*^−/−^ *Ucp1*^−/−^ mice during 6 days of CysF diet (n=7 *Cth*^−/−^ and n=8 *Cth*^−/−^ *Ucp1*^−/−^). m-n) RNA-seq based expression of genes associated with m) creatine futile cycle (*Slc6a8*, *Gatm*, *Gamt*, *Ckmt2*, *Alpl* and *Ckb*) and n) calcium futile cycle (*Atp2a2* and *Ryr2*) in the SFAT of *Cth*^+/+^ and *Cth*^−/−^ mice fed CTRL or CysF diet for 6 days (n=4/group). o) qPCR gene expression of *Sarcolipin* and *Atp2a2* in the soleus of *Cth*^+/+^ and *Cth*^−/−^ mice fed CTRL or CysF diet for 6 days (n=3 *Cth*^+/+^CTRL, n=6 *Cth*^+/+^CysF, n=3 *Cth*^+/+^CTRL and n=5 *Cth*^+/+^CysF). p) RNA-seq based expression of genes associated with triglyceride and fatty acid metabolism (*Dgat1*, *Pnpla2*, *Lipe*, *Gk*) in the SFAT of *Cth*^+/+^ and *Cth*^−/−^ mice fed CTRL or CysF diet for 6 days (n=4/group). Data are expressed as mean±SEM. Statistical differences were calculated by 2-way ANOVA with Sidak’s correction for multiple comparisons, and by unpaired t-test (*p<0.05, **p<0.01, ***p<0.001).

**Extended Data Figure 6:**
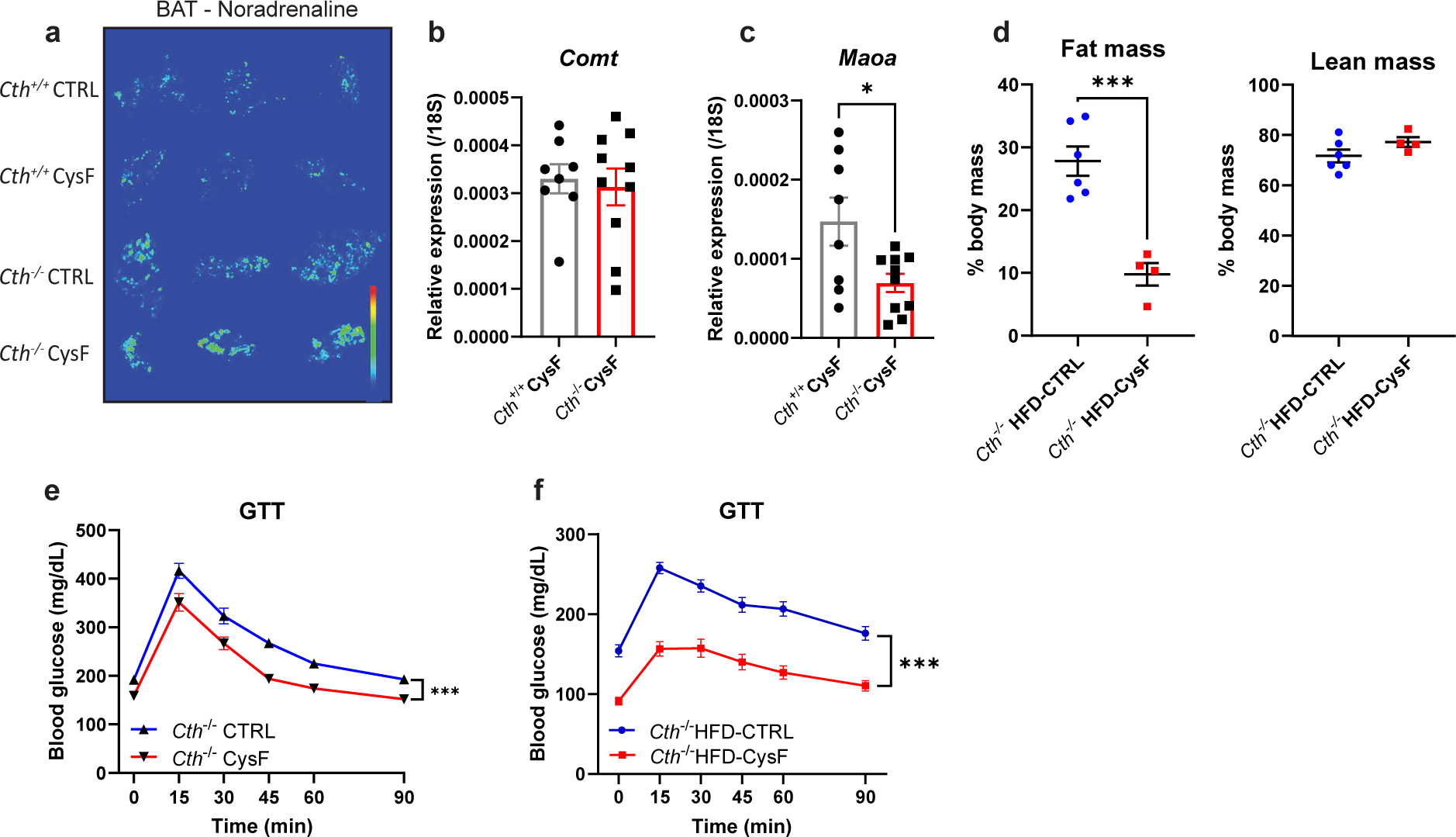
Cysteine starvation induced browning requires adrenergic signaling. a) Imaging mass spectrometry of noradrenaline in the BAT of *Cth*^+/+^ and *Cth*^−/−^ fed 6 days of CTRL or CysF diet. b-c) qPCR gene expression of b) *Maoa* and c) *Comt* in SFAT of *Cth*^+/+^ (n=8) and *Cth*^−/−^ (n=10) mice fed with CysF diet for 6 days. d) Body composition measured by Echo-MRI on day 6 post diet switch (n=6 *Cth*^−/−^ HFD-CTRL and n=4 *Cth*^−/−^ HFD-CysF). e) The glucose tolerance test (GTT) in mice fed control and cysF diet with glucose dose based on lean mass. f) The GTT in *Cth*^−/−^ after diet switch from HFD-CTRL to HFD-CysF (*Cth*^−/−^ HFD-CTRL n=19, *Cth*^−/−^ HFD-CysF, n=20). The glucose administration based on total body-weight. Data are expressed as mean±SEM. Statistical differences were calculated by 2-way ANOVA with Sidak’s correction for multiple comparisons, and by unpaired t-test (*p<0.05, ***p<0.001).

